# Molecular basis for the transcriptional regulation of an epoxide-based virulence circuit in *Pseudomonas aeruginosa*

**DOI:** 10.1101/2024.01.16.572601

**Authors:** Susu He, Noor M. Taher, Kelli L. Hvorecny, Michael J. Ragusa, Christopher D. Bahl, Alison B. Hickman, Fred Dyda, Dean R. Madden

**Affiliations:** Department of Biochemistry and Cell Biology, Geisel School of Medicine at Dartmouth, Hanover, NH 03755 USA; Department of Chemistry, Dartmouth, Hanover, NH 03755 USA; Laboratory of Molecular Biology, NIDDK, National Institutes of Health, Bethesda, MD 20892 USA

**Keywords:** *Pseudomonas aeruginosa*, Epoxide-based virulence circuit, TetR family transcriptional regulators (TFRs), X-ray crystallography, Protein-DNA recognition

## Abstract

The opportunistic pathogen *Pseudomonas aeruginosa* infects cystic fibrosis (CF) patient airways and produces a virulence factor Cif that is associated with worse outcomes. Cif is an epoxide hydrolase that reduces cell-surface abundance of the cystic fibrosis transmembrane conductance regulator (CFTR) and sabotages pro-resolving signals. Its expression is regulated by a divergently transcribed TetR family transcriptional repressor. CifR represents the first reported epoxide-sensing bacterial transcriptional regulator, but neither its interaction with cognate operator sequences nor the mechanism of activation has been investigated. Using biochemical and structural approaches, we uncovered the molecular mechanisms controlling this complex virulence operon. We present here the first molecular structures of CifR alone and in complex with operator DNA, resolved in a single crystal lattice. Significant conformational changes between these two structures suggest how CifR regulates the expression of the virulence gene *cif*. Interactions between the N-terminal extension of CifR with the DNA minor groove of the operator play a significant role in the operator recognition of CifR. We also determined that cysteine residue Cys107 is critical for epoxide sensing and DNA release. These results offer new insights into the stereochemical regulation of an epoxide-based virulence circuit in a critically important clinical pathogen.

## INTRODUCTION

*Pseudomonas aeruginosa* is an opportunistic pathogen that infects immunocompromised patients. It can cause acute and chronic airway infections, particularly in patients with underlying lung disease. For example, *P. aeruginosa* chronically colonizes the majority of adult patients with cystic fibrosis (CF) and is a major contributor to respiratory failure in these patients (1). While infections by *P. aeruginosa* are not as frequent in patients with chronic obstructive pulmonary disease (COPD) or non-CF bronchiectasis, these infections are associated with high rates of mortality (2–5). The success of *P. aeruginosa* in establishing persistent lung infections reflects the fact that it harbors diverse mechanisms of genetic adaptation and antibiotic resistance (1). For example, the bacteria secrete virulence factors that manipulate host physiology, subvert host defenses, and favor colonization (1, 6).

Previous studies have uncovered a mechanistically distinct secreted virulence factor from *P. aeruginosa* that can reduce the cell-surface abundance of the cystic fibrosis transmembrane conductance regulator (CFTR) in airway epithelial cells (6) and degrade pro-resolving signals typically formed during the response to airway infection (7). The virulence factor responsible for these effects, the CFTR inhibitory factor (Cif), is also capable of reversing pharmacological rescue of the most common disease-associated CFTR mutation in human airway epithelial cells (8). Cif is an epoxide hydrolase, an enzyme that converts an oxirane ring into a vicinal diol (9, 10), and its hydrolase activity is required for its effects on CFTR (11). Epoxides are cyclic ethers with a three-atom ring and play important physiological roles (7, 12, 13). Within cells, Cif accelerates degradation of CFTR by inhibiting its post-endocytic deubiquitination (14). Cif thus promotes bacterial colonization, as demonstrated in a mouse model of acute pneumonia (15). Cif also hydrolyzes the epoxy-polyunsaturated fatty acid 14,15-epoxyeicosatrienoic acid (14,15-EET), a signal required for the formation of the pro-resolving mediator 15-epi-lipoxin A_4_. Consistent with these mechanistic observations, higher levels of Cif protein in pediatric airway samples correlated with worse lung function and increased hyperinflammatory signaling (7).

Exposure to epibromohydrin (EBH) and other synthetic epoxides potently upregulates *cif* expression in *P. aeruginosa*, suggesting that it is part of an epoxide-responsive circuit (16). Indeed, the sequence of Cif is homologous to a protein that was earlier found to enable a soil-dwelling pseudomonad to degrade environmental epichlorohydrin (17). *cif* is part of an operon that is adjacent to a divergently transcribed gene *cifR* (Figure 1A; top row), which exhibits sequence homology to TetR family transcriptional regulators (TFRs). Expression of the *cif* gene is repressed by CifR binding at the promoter immediately upstream of the *cif* operon (Figure 1A; middle row), and clinical strains exhibiting high *cif* expression often show reduced levels of *cifR* expression (16). Furthermore, EBH disrupts CifR binding to the intergenic promoter region, permitting transcription of *cifR* and the three divergently transcribed genes including *cif* (16, 18) (Figure 1A; bottom row). Cif-mediated hydrolysis of the epoxide (Figure 1A; bottom right) has the potential to reset the transcriptional state. Thus, based on sequence and functional analysis, CifR appears to be a TetR repressor and to represent the first reported epoxide-sensitive bacterial transcriptional regulator.

**Figure 1.**
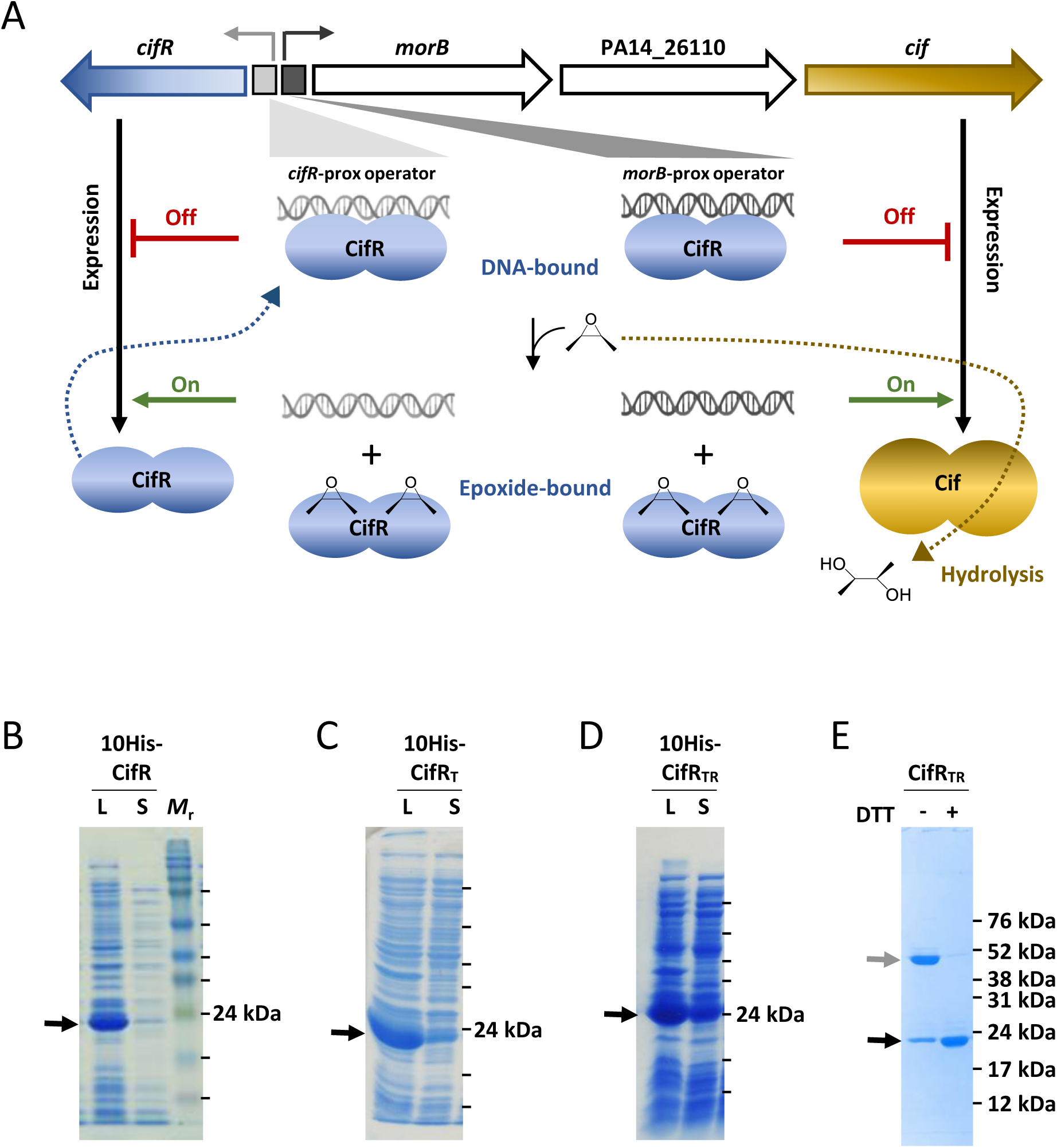
The *cif*-*cifR* operon and CifR protein solubility. **(A)** *Top row:* The *cif*-*cifR* operon. The arrows show the four open reading frames in the operon. The two rectangles in grey represent the two operator sites for CifR binding (*Middle row*; blue). *Bottom row*: Binding to an epoxide (schematic drawing) releases CifR from the DNA. Cif is shown as a gold oval doublet which converts the epoxide ring to a vicinal diol (schematic drawing). **(B-D)** Coomassie-stained SDS-PAGE gels for CifR constructs. Rosetta 2 (DE3) cells were transformed with plasmids expressing decahistidine-tagged WT (B), CifR_T_ (C), and CifR_TR_ (D) constructs, grown in LB medium, and induced with 0.5 mM IPTG overnight at 16 °C. Lanes are marked “L” for whole-cell lysate or “S” for supernatant clarified by centrifugation at 40,000 rpm for 1 hr in a 45 Ti rotor. The positions of *M*_r_ standards are shown for each gel (adjacent lines) and the corresponding *M*_r_ values shown in (E). The short black lines on the side of the gel indicates protein standards of size 76 kDa, 52 kDa, 38 kDa, 31 kDa, 24 kDa, 17 kDa and 12 kDa from top to bottom. The black arrows point to the expected gel migration position for CifR monomer according to the protein standards. The grey arrow points to the expected gel migration position for CifR dimer. **(E)** SDS-PAGE gel of purified CifR_TR_ (in the absence and presence of reducing agent) following cleavage of the decahistidine tag.

To resolve the molecular basis for CifR-operator recognition and lay a foundation for understanding the epoxide responsiveness, we solved the X-ray crystal structure of CifR in complex with double-stranded (ds) operator DNA at 2.6 Å resolution. Despite the length of the operator sequence, the overall structure turned out to involve a single CifR dimer binding per operator site, with unusual protein-DNA contacts at the periphery. Surprisingly, the lattice of the selenomethionine derivative used for phase determination was found to harbor a DNA-free CifR molecule caged within a bulk-solvent channel. Thus, we were able to determine both the DNA-bound and DNA-free structures of CifR from a single lattice form, revealing conformational changes associated with DNA binding and suggesting a hypothesis for epoxide-mediated release of DNA relying on a critical cysteine residue.

## MATERIAL AND METHODS

### Strains and plasmids

*Escherichia coli* DH5α cells were used for cloning. BL21-derived Rosetta 2 (DE3) cells were used for protein expression. The *cifR* gene (PA14_26140; UniProt ID: A0A0H2ZCS5) and its derivative mutants were cloned in plasmid pET-16b with a 10-His tag followed by a human rhinovirus 3C protease cleavage site between the tag and the N-terminus of the recombinant proteins (19). Mutagenesis of *cifR* was performed using the Pfu Turbo DNA Polymerase (Agilent), and mutants were verified by Sanger sequencing. Primers used for cloning are listed in Table S1.

### Protein expression and purification

Cells transformed with the relevant expression plasmid (either from freshly transformed cells or from a glycerol stock) were first grown as a starter culture in 50 mL of Lysogeny Broth (LB) with 100 µg/mL ampicillin and 34 µg/mL chloramphenicol at 37 °C in Erlenmeyer flasks overnight. Ten milliliters of this starter culture were inoculated into 1 L LB with ampicillin and chloramphenicol and cultivated at 37 °C in 2 L Erlenmeyer flasks. Protein expression was induced when OD_600_ reached a value of 0.4-0.6 by addition of isopropyl β-D-1-thiogalactopyranoside (IPTG) to a final concentration of 0.5 mM and was allowed to continue overnight at 16 °C. Cells were collected by centrifugation, resuspended in 25 mM 4-(2-hydroxyethyl)-1-piperazineethanesulfonic acid (HEPES) pH 7.5, 1 M NaCl, 5 mM β-mercaptoethanol, and 10 mM imidazole pH 8.5, and then lysed using a microfluidizer (Model M-110L, Microfluidics, US) equipped with an H10Z interaction chamber submerged in ice. Following sequence optimization (see Results for details), CifR_TR_ was subsequently purified by immobilized metal affinity chromatography (IMAC) using HisPur Ni-NTA agarose resin (Thermo Fisher Scientific) in 25 mM HEPES pH 7.5, 1 M NaCl, and 5 mM β-mercaptoethanol, supplemented with varying concentrations of imidazole using a step-elution protocol. The imidazole concentration was 10 mM in the binding buffer, 100 mM in the washing buffer and 500 mM in the elution buffer. Two hundred microliters of 4 mg/mL hexahistidine-tagged 3C protease were added to the 40 mL eluted fraction pool. The mixture was dialyzed against 2 L of 25 mM HEPES pH 7.5, 200 mM NaCl, 1 mM dl-dithiothreitol (DTT), and 40 mM imidazole pH 8.5 overnight at 4 °C. This sample was reapplied to Ni-NTA agarose resin to capture any uncleaved protein along with the 3C protease. The flow-through fraction was concentrated and centrifuged to remove any precipitated protein. The supernatant was subjected to size-exclusion chromatography (SEC) at 4°C using a Superdex 75 26/600 column equilibrated with Buffer A (25 mM HEPES, pH 7.5, 200 mM NaCl, and 1 mM DTT).

To obtain selenomethionine (SeMet) labeled protein, we inhibited the methionine (Met) biosynthesis pathway by adding high concentrations of isoleucine, lysine, and threonine (20). Starter cultures were grown in Terrific Broth (TB) supplemented with appropriate antibiotics at 37 °C, until the OD_600_ reached a value of 3-4. Cells were pelleted at 2,000 × *g* for 10 min, resuspended in modified M9 minimal medium (21). The culture was incubated at 37 °C for 1 h to minimize the amount of residual methionine. Amino acids were added to the culture to reach final concentrations of 100 µg/mL each of selenomethionine, lysine, phenylalanine, and threonine, and 50 µg/mL each of isoleucine, leucine, and valine. Cells were transferred to an incubator at 20 °C, incubated for a further 30 min, and then protein expression was induced by addition of 0.5 mM IPTG, followed by incubation at 20 °C overnight.

### Electrophoretic mobility shift assay (EMSA)

Protein-DNA binding reactions were carried out in 10 µL reaction volumes containing varying concentrations of DNA and protein (see figures for details) in 25 mM HEPES, pH 7.5, 150 mM NaCl and 0.5 mM DTT. EBH and *N*-ethylmaleimide (NEM) were initially dissolved in dimethyl sulfoxide (DMSO) at 100 mM. Serial dilutions were prepared with Buffer A, and one microliter of each dilution of EBH or NEM was added to an aliquot of the protein-DNA mix. The reaction was incubated at RT for 30 min. Three microliters of loading dye (50% [*w/v*] glycerol, 0.04% [*w/v*] cyanole xylene) were added, and the samples were loaded on a 12% Tris/borate/EDTA native gel. Electrophoresis was performed for 130 min at 80 V to separate protein-bound DNA and free DNA. The gel was stained with SYBR Gold Nucleic Acid Gel Stain (Invitrogen) and imaged using a Gel Doc™ XR+ Gel Documentation System (Biorad).

### Crystallization and data collection

The concentrations of the protein dimer and DNA duplex in crystallization screens were 100 µM and 125 µM, respectively. Initial screening was carried out for CifR mixed with DNA fragments varying from 23bp to 36bp using at least 10 commercially available screens. Crystals were obtained under more than 50 conditions and were assayed for diffraction quality. The conditions that yielded the best diffraction were further optimized by varying pH and precipitant concentration. For the CifR_TR_:dsDNA_26L_ native co-crystal, the best dataset was obtained from a crystal grown in 0.1 M Tris pH 8.3, 0.2 M MgCl_2_, and 16% [*w/v*] polyethylene glycol (PEG) 4000, conditions that were also used to obtain CifR_TR_^C107S^:dsDNA_26L_ co-crystals. For the CifR_TR_:dsDNA_26L_ SeMet co-crystal, the best condition was 0.1 M Tris pH 7.5, 0.2 M MgCl_2_, 10% [*w/v*] PEG 1000, 10% [*w/v*] PEG 8000. The final SeMet and native datasets of CifR_TR_:dsDNA_26_ used for structure determination were collected at NSLS-II beamline 17-ID-1. Diffraction data were collected on an Eiger 9M detector through 360° of total rotation using the NSLS-II AMX beamline’s vector collection mode (22) in 0.2° frames at a detector distance of 200 mm. For the SAD experiment using the SeMet crystals, the optimal peak energies were determined from X-ray absorption scans. The native dataset for the CifR_TR_^C107S^:dsDNA_26L_ co-crystal was collected at NSLS-II beamline 17-ID-2 on an Eiger 16M detector through 360° of total rotation using vector collection mode in 0.2° frames at a detector distance of 300 mm. See Table 1 for additional details.

**Table 1:**
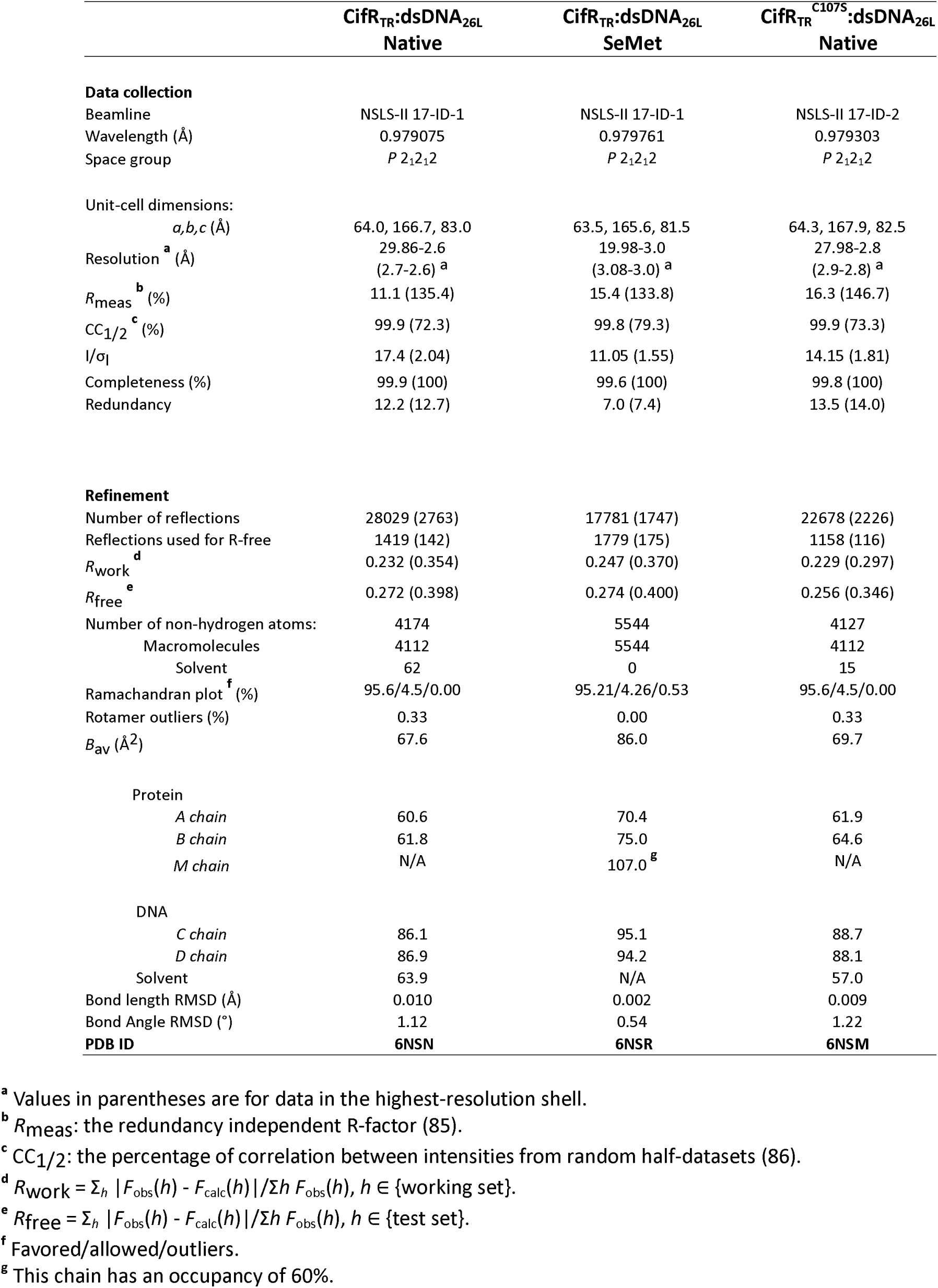
Data collection and refinement statistics.

### Diffraction data processing and structure refinement

Datasets were indexed, integrated, and scaled internally using the XDS package (23). Experimental phases were determined using the SeMet dataset with PHENIX_Autosol (24) and were used as the basis for initial model building of a CifR_TR_ dimer bound to a dsDNA_26L_ duplex. The structure was refined by iterative rounds of manual fitting using COOT (25) alternating with rigid-body, conjugate-gradient, real-space, and/or individual B-factor refinement using PHENIX_Refine (24). As described below (Results), during this process, an additional, DNA-free (apo-)CifR monomer (Chain M) was identified in the asymmetric unit, modeled, and refined. Grouped occupancy refinement was performed for chain M (apo-CifR) to estimate occupancy. The asymmetric unit contained 3 CifR monomers and one 26 bp DNA duplex.

Given differences in unit cells, initial phases for the higher-resolution native datasets were obtained by molecular replacement using the experimentally phased and refined CifR_TR_:dsDNA_26L_ structure as a starting model. The apo-CifR could not be modeled in the native dataset, due to substantially weaker electron density (Figure S4B). For structural comparisons, we used the higher-resolution native structures of the chains A and B (DNA-bound CifR_TR_ dimer) together with the SeMet structure of chain M (apo CifR). The N-terminal two residues of Chain A (native dataset), four residues of Chain B (native), and nine residues of Chain M (SeMet) could not be modeled due to disordered electron density. The PyMol Molecular Graphics System (Schrödinger, LLC) was used for least-squares superpositions and for visualization of structures and of difference and Polder (omit) electron-density maps (26). Electrostatic surface potential was calculated using the APBS plugin.

### Analytical ultracentrifugation (AUC)

Sedimentation velocity was determined using a Beckman Coulter ProteomeLab XL-A ultracentrifuge. The reference cell chamber was loaded with 410 µl of Buffer A. The sample chamber was loaded with 400 µL of either protein, DNA, or a protein-DNA mixture. The amount of sample was adjusted to yield an absorbance at 280 nm (A_280_) of approximately 0.4. In the protein-DNA mixture, the DNA concentration was chosen to yield a 1.25-fold molar excess relative to the concentration of protein dimers. The samples were spun at 35,000 rpm in an An-60 Ti rotor at 20 °C with reads every 3 min for at least 200 scans. Data from the first 100 scans were analyzed.

### Calculation of shape-independent macromolecular MW

SEC partition coefficients (K_av_) were determined for CifR_TR_, dsDNA, and the CifR_TR_-dsDNA complex. Given the elongated shape of DNA duplexes, instead of calibrating K_av_ directly vs MW, we used the Svedberg and Stokes-Einstein equations to estimate the MW of each macromolecule in this study according to Equation 1:

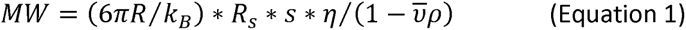

In brief, we calibrated K_av_ vs. Stokes radii (*R*_S_) for standard proteins, plotting (-logK_av_)^1/2^ vs. *R*_S_ (27). The sedimentation coefficients (*s*) were obtained by fitting a continuous c(*s*) distribution to experimental AUC traces using SEDFIT (28). The buffer density ρ (1.0084 g/mL) and viscosity η (0.010386 g/cm/sec) were calculated using SEDNTERP (29). The partial specific volume *v̄* (0.73075 mL/g) for CifR_TR_ was calculated from the protein sequence using SEDNTERP. The partial specific volume for DNA was set at 0.55 mL/g (30). The partial specific volume for protein-DNA complex was calculated as the weight average of the partial specific volumes of a DNA duplex (31) with either one or two CifR_TR_ dimers bound per dsDNA_26L_ duplex.

### Small-angle X-ray scattering (SAXS)

Protein-DNA mixtures were prepared in Buffer A, with protein concentrations of 3 mg/mL, 2 mg/mL, or 1 mg/mL, together with DNA duplex at concentrations equivalent to a 1:1 dimer:DNA molar ratio for each mixture. The mixture was subjected to SEC using a Superdex 200 10/300 column to confirm the formation of protein-DNA complex. A sample of isolated DNA duplex was prepared at a concentration of 325 µM. The samples were measured at the LiX beamline (32) at the National Synchrotron Light Source II (NSLS-II), using the standard conditions for protein solution scattering. For each sample, the scattering data were averaged from three 3-second exposures. Buffer subtraction was based on the water peak intensity (33). Data were analyzed over a q range of 0.02 to 0.25 Å^-1^ using PRIMUS (34). Guinier plots were found to be linear, consistent with minimal levels of protein aggregation, and were used to determine the radius of gyration (R_g_) for each sample. Across varying sample concentrations, R_g_ values were consistent within 0.1 Å, suggesting that neither the overall structure nor the oligomeric state are strongly dependent on concentration in this range. Theoretical SAXS curves were calculated from crystal structures using the FoXS webserver (35, 36). Ten independent molecular envelopes were calculated from the 3 mg/ml SAXS data using DAMMIF, averaged using DAMAVER and then refined using DAMMIN (37–39). PDB structures were aligned with the molecular envelope using SUPCOMB (40).

## RESULTS

### Mutagenetic stabilization of the CifR protein

To investigate the regulation of CifR at the molecular level, we needed to address the lack of solubility of recombinantly expressed 10His-CifR (19). Most of the wild-type protein pelleted during high-speed centrifugation (Figure 1B), and the protein that could be purified continued to aggregate. We suspected that protein solubility could be improved by decreasing the number of cysteine residues in the construct. Sequence similarity with other TFRs suggests that CifR is composed of a relatively conserved N-terminal DNA-binding domain with a typical HTH motif and a more variable C-terminal ligand-binding domain, which contains all three of the cysteine residues found in CifR (Figure S1). Cys107 is found in 52% of 938 TFR sequences analyzed, whereas Cys99 and Cys181 are each found in less than 3% (Figure S1). We therefore investigated replacement of Cys99 and Cys181 with threonine and arginine, which are respectively the most prevalent residues at their positions in the TFR sequence alignment.

While a C99T mutation alone did not improve initial solubility, the double-cysteine mutations (C99T/C181R; or CifR_TR_) substantially increased the fraction of protein remaining in the clarified lysates (Figure 1B-D). Immobilized metal-affinity and size-exclusion chromatography yielded pure protein with a size corresponding to a CifR_TR_ monomer as determined by SDS-PAGE (Figure 1E). The SEC elution profile of CifR_TR_ was consistent with the formation of a protein dimer in the absence of DNA (with a calibrated molecular mass of 43 kDa; Figure S2). The dsDNA_36_ forms a protein-DNA complex with CifR_TR_ as confirmed by EMSA (Figure 2B, left-hand panel) and SEC (Figure S2A). It is worth noting that apo-CifR_TR_ is still prone to crosslinking as a dimer in the absence of reducing agent (Figure 1E), with slow precipitation occurring over time. Interestingly, a mixture of CifR_TR_ and dsDNA_36_ did not show visible precipitation, indicating the presence of its operator DNA could stabilize this protein. Based on this observation, we sought to obtain stable protein-DNA complex, which required DNA optimization.

**Figure 2.**
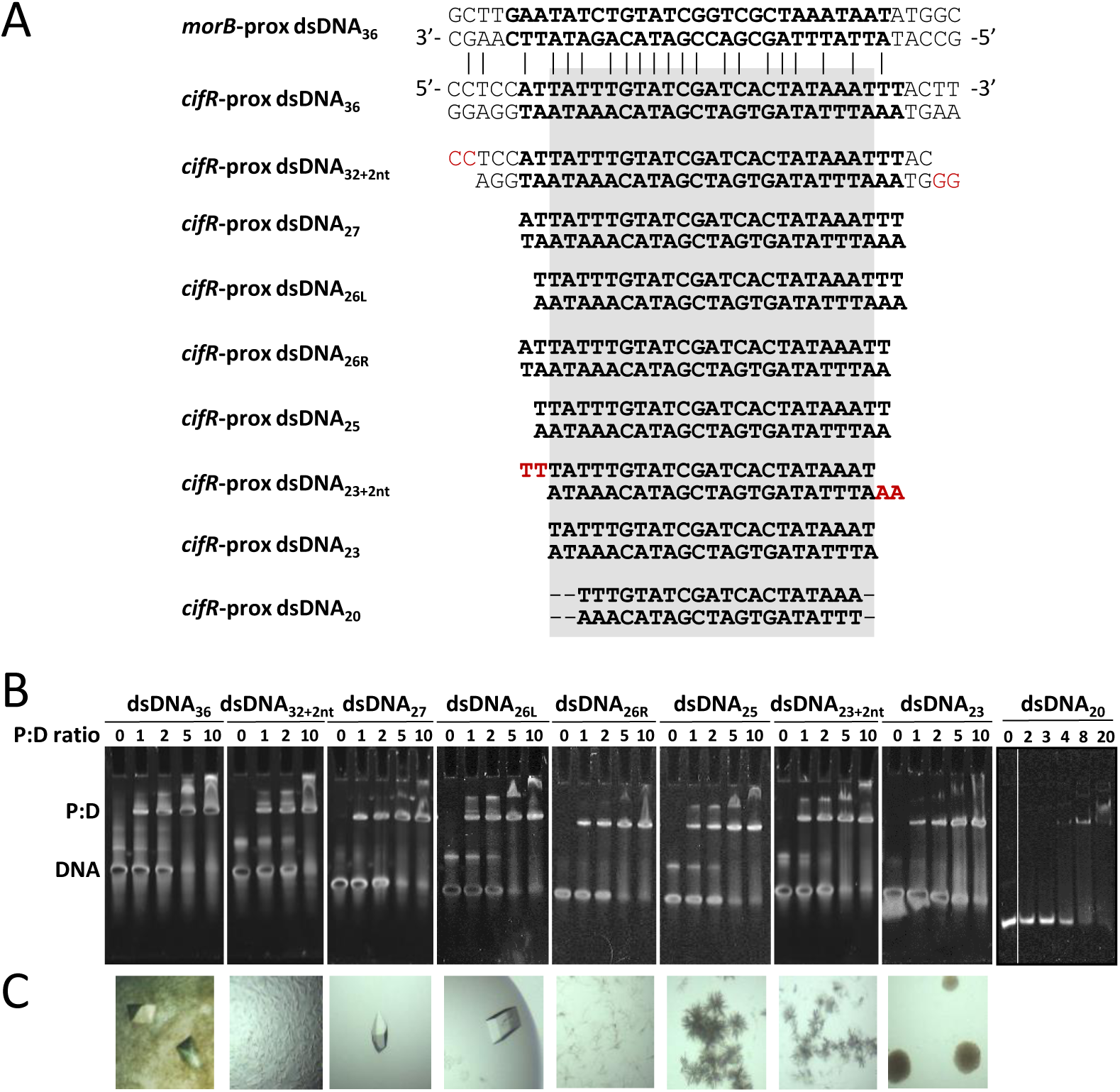
DNA screening for CifR_TR_:DNA complex formation and crystallization. **(A)** Sequences of the DNA oligonucleotides tested for optimizing the CifR_TR_:DNA complex formation. Varying sequence lengths of the *cifR*-proximal operator DNA (*2^nd^ row*) were assessed for CifR_TR_:DNA complex formation and co-crystallization. The sequence highlighted in grey represents the minimal length of DNA (23bp) which could bind to CifR in this study. Cohesive bases are highlighted in red. Sequence alignment of the cifR-proximal and the morB-proximal operator (*Top row*) is shown. The two operators present in an inverted orientation in the genome of *P. aeruginosa*. The orientation of the strands labeled with 5’ to 3’ correspond to the top strand in Figure 1A. **(B)** EMSAs of CifR_TR_ with each of the oligonucleotides listed in (A). The concentration of DNA in the assay was 2 µM. “P:D ratio” is the molar ratio of CifR monomer to DNA. “P:D” and “DNA” indicate the migration positions of protein:DNA complex and free DNA, respectively. This shorthand is used throughout the article. **(C)** Representative crystallization screening results for each complex assayed in (B). Each picture shows the crystal morphology with the best diffraction, or the most common result observed during the screening process.

### Operator “footprinting” and crystallization of the CifR_TR_:operator complex

Our previous studies had identified two distinct operator DNA sites for CifR in the *P. aeruginosa* genome (Figure 1A and Figure 2A) (16). Both sites are located in the intergenic region between *cifR* and *morB*, the gene located at the 5’ end of the *cif* operon (16). For each site, full CifR:DNA binding affinity was observed with a 27 bp construct (Figure 2A, sequence in bold) and lost with a 16-25 bp construct (16). We tested a series of dsDNA constructs varying from 36 bp to 20 bp in length. We only focused on the operator DNA site which is proximal to *cifR* since this site showed 10-fold higher binding affinity compared to the one proximal to *morB* (16). We also tested two DNAs with cohesive 2 bp overhangs, designed to stabilize potential lattice interactions. We measured the binding affinity of each of these DNA constructs (Figure 2A and Table S1) for CifR_TR_ by EMSA and found that CifR_TR_ could bind to DNAs with length varying between 36 bp and 23 bp with comparable affinities (Figure 2B). Further shortening the DNA to 20 bp progressively hindered the formation of a robust complex at single micromolar concentrations (Figure 2B). Based on these results, we concluded that dsDNA_23_ contains the minimum binding-site footprint for CifR, and we took 23 bp as the lower boundary on the length of dsDNA for crystallization studies.

Initial vapor-diffusion crystallization screening trials were set up with distinct mixtures of CifR_TR_ with each of the different lengths of DNA that form complexes (Figure 2B) at a 1.25-fold molar excess of DNA. Crystals of varying quality were obtained with most of the constructs tested, as shown in Figure 2C. Complexes with the dsDNA_36_, dsDNA_27_, and dsDNA_26L_ constructs produced crystals with good size and morphology, with crystallization conditions shown in Table S2. The crystals obtained with the dsDNA_26L_ construct showed good overall quality and yielded a native dataset at 2.6 Å resolution (Table 1).

### Co-crystallization of two protein conformations: CifR_TR_:dsDNA_26L_ complex and apo-CifR_TR_

We initially attempted to solve the structure using molecular replacement (MR) with a native dataset. In all known cases, TFRs form homodimeric complexes in the absence of DNA (41–45). The structures of TFR:DNA complexes reveal two fundamentally distinct stoichiometries, with a single operator binding either one or two TFR dimers (Figure S3). The Matthews coefficient (46) was consistent with a single DNA duplex bound to either one (∼67% solvent) or two (∼41% solvent) CifR dimers in the ASU. However, search models based on the published structures of other TFRs (Figure S3) failed to return acceptable solutions in either stoichiometry. Therefore, we used experimental phasing by SeMet substitution. Following expression and purification through IMAC and SEC, an average labeling efficiency of ∼76% was determined by MALDI-TOF. Optimized SeMet crystals diffracted to 3.0 Å resolution. Electron-density maps based on SeMet-SAD phasing were generated in Phenix and used for initial model building and refinement, revealing a single CifR_TR_ dimer bound to each dsDNA_26L_ duplex, consistent with SEC results for this complex (Figure S2B).

The crystal lattice is formed in one orientation by head-to-tail contacts between DNA molecules, forming extended helices (Figure 3A). CifR_TR_-dsDNA_26L_ complexes are oriented alternately up and down along the extended DNA helices (Figure 3A, bottom strand). The resulting rows of protein:DNA complexes pack at right angles to extend the lattice in a second dimension, contacting each other through protein:protein interactions (Figure 3A, right panel). Consistent with the relatively high (∼67%) solvent content calculated for a single CifR_TR_-dsDNA_26L_ complex per ASU, this arrangement creates relatively large channels that run the length of the crystal lattice (Figure 3B) and that span paired ASUs related by two-fold axes associated with the *P* 2_1_2_1_2 space group. These channels contained weak, but consistent experimental electron density, as well as a subset of the original candidate heavy-atom sites on either side of the two-fold axis that were not accounted for by the CifR_TR_-dsDNA_26L_ complex (Figure S4A). We were able to model this additional density as a DNA-free (apo-)CifR_TR_ dimer that was apparently trapped between adjacent CifR_TR_-dsDNA_26L_ complexes (Figure 3B, right) and aligned along a crystallographic symmetry axis. The asymmetric unit (ASU) in the SeMet lattice is thus composed of a CifR_TR_ dimer bound to dsDNA_26L_ together with an additional apo-CifR_TR_ monomer, which forms a dimer with its symmetry mate (PDB ID: 6NSR). The solvent content of this hybrid ASU was estimated at 54%. It is important to note that the overall electron density for the “stowaway” apo-CifR_TR_ protein remained substantially weaker than that of CifR_TR_-dsDNA_26L_ complex, even after model building and refinement. This is reflected in the refined parameters of the apo-CifR_TR_ model (Chain M), which yielded an occupancy of only 60% and an average B-factor higher than either of the DNA-bound monomers (chains A and B; Table 1).

**Figure 3.**
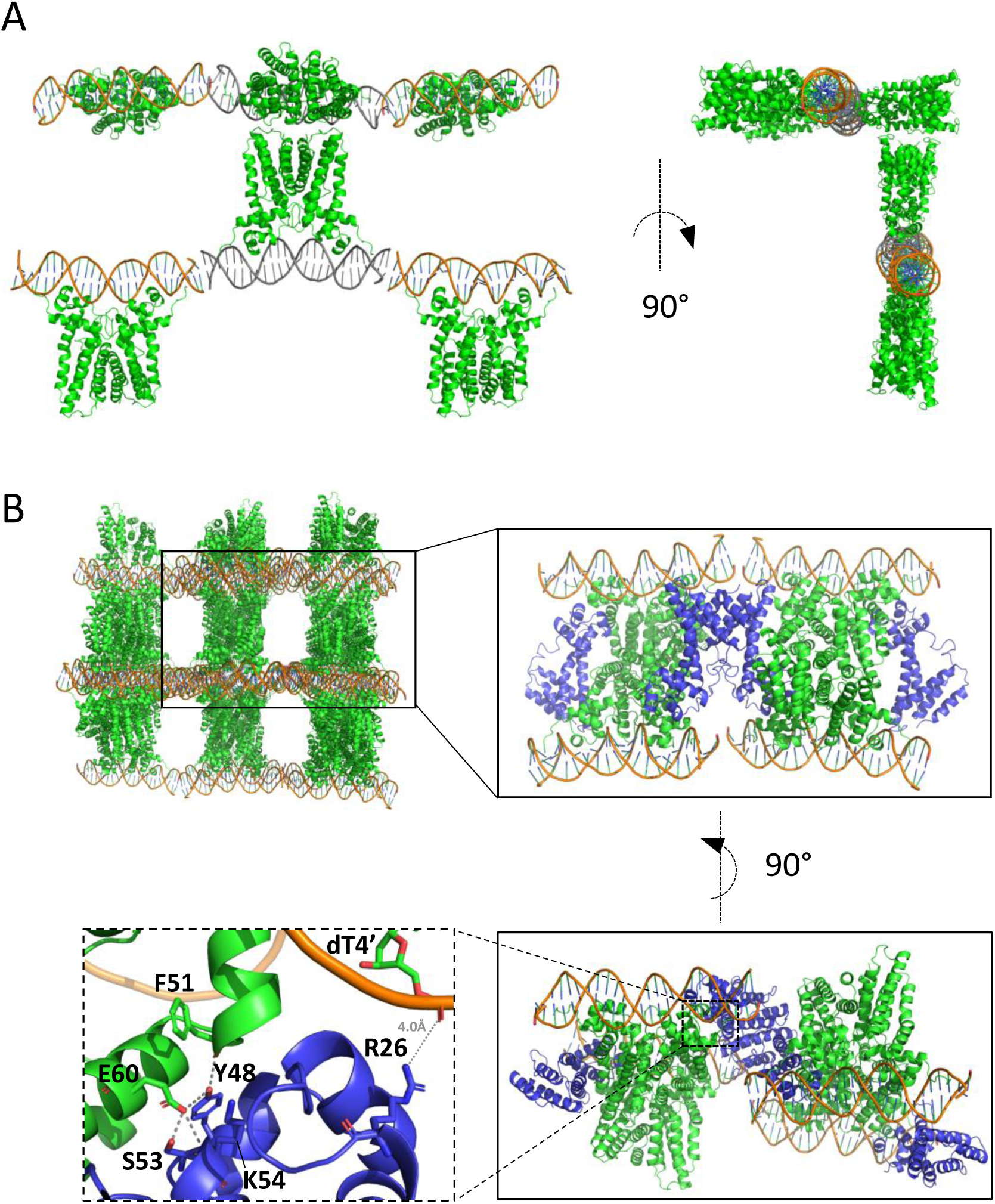
Lattice packing of the CifR_TR_:dsDNA_26L_ complex and apo-CifR_TR_ crystal. **(A)** Lattice formation by DNA- and protein-protein interactions. DNA duplexes (orange/grey) align end-to-end to form extended helices, with each DNA molecule bound by one CifR_TR_ dimer (green ribbons). The DNA duplexes in the center are shown in grey to distinguish the DNA-DNA junctions. A 90° rotation about the indicated axis reveals the protein-protein contacts that join neighboring DNA helical extensions in the lattice (right). **(B)** Apo-CifR_TR_ is caged into the bulk solvent channel of the lattice formed by CifR_TR_:dsDNA_26L_. The panel on the top left shows the bulk solvent channels formed by the CifR_TR_:dsDNA_26L_ complex. The inset on the top right shows a closeup view of the apo-CifR_TR_ dimers (blue ribbons), which are omitted in the parent panel. The inset on the bottom right shows the 90° rotation view of the inset on the top right. The inset on the bottom left shows the close up view of lattice contacts between the apo-Cif_TR_ and CifR_TR_:dsDNA_26L_ complex. Residues with polar contacts are shown as sticks.

No direct interactions were observed between this apo-CifR_TR_ and either of the DNA duplexes, even though from one perspective (Figure 3B, 90° rotation), this apo-CifR_TR_ dimer appears to bridge DNA duplexes in the lattice. In fact, the closest apo-CifR_TR_ residue is Arg26 from helix α1, the nitrogen (NH_2_) group of which is 4.0 Å away from the oxygen group (OP1) of the DNA backbone (dT4’). Instead, apo-CifR_TR_ interacts with adjacent protein molecules: direct polar contacts were observed between apo-CifR_TR_ and the CifR_TR_ protein in the CifR_TR_:dsDNA_26L_ complex (Figure 3B, bottom left). Tyr48 from helix α3 of apo-CifR_TR_ forms a hydrogen bond with the main chain of Phe51, which is at the loop between helices α3 and α4 in the CifR_TR_:dsDNA_26L_ complex. This Tyr48, together with Ser53 and the main-chain amide of Lys54 from helix α4 of apo-CifR_TR_, form polar contacts with Glu60 from helix α4 of CifR_TR_:dsDNA_26L_ complex (Figure 3B, bottom left).

To determine if the crystal structure of the CifR_TR_:dsDNA_26L_ complex represents the major component of the solution state, theoretical SAXS curves were calculated and were found to fit very well to the experimental SAXS data from a 1:1 molar mixture of the two components (χ^2^=0.39; Figure S5A). In addition, the CifR_TR_:dsDNA_26L_ complex structure fits into the molecular envelope calculated from the SAXS data with little additional density (Figure S5B). SEC and AUC also provide shape-independent molecular mass estimation by applying Svedberg and Stokes-Einstein equations (see eqn 1), yielding results consistent with the observed CifR_TR_:dsDNA_26L_ complex (47) (Table 2, Figures S2 and S6). These results together confirm that the crystal structure of the CifR_TR_:dsDNA_26L_ complex represents the predominant form of the complex in solution, for both the 26bp and the full-length 36bp operator sequences.

**Table 2:**
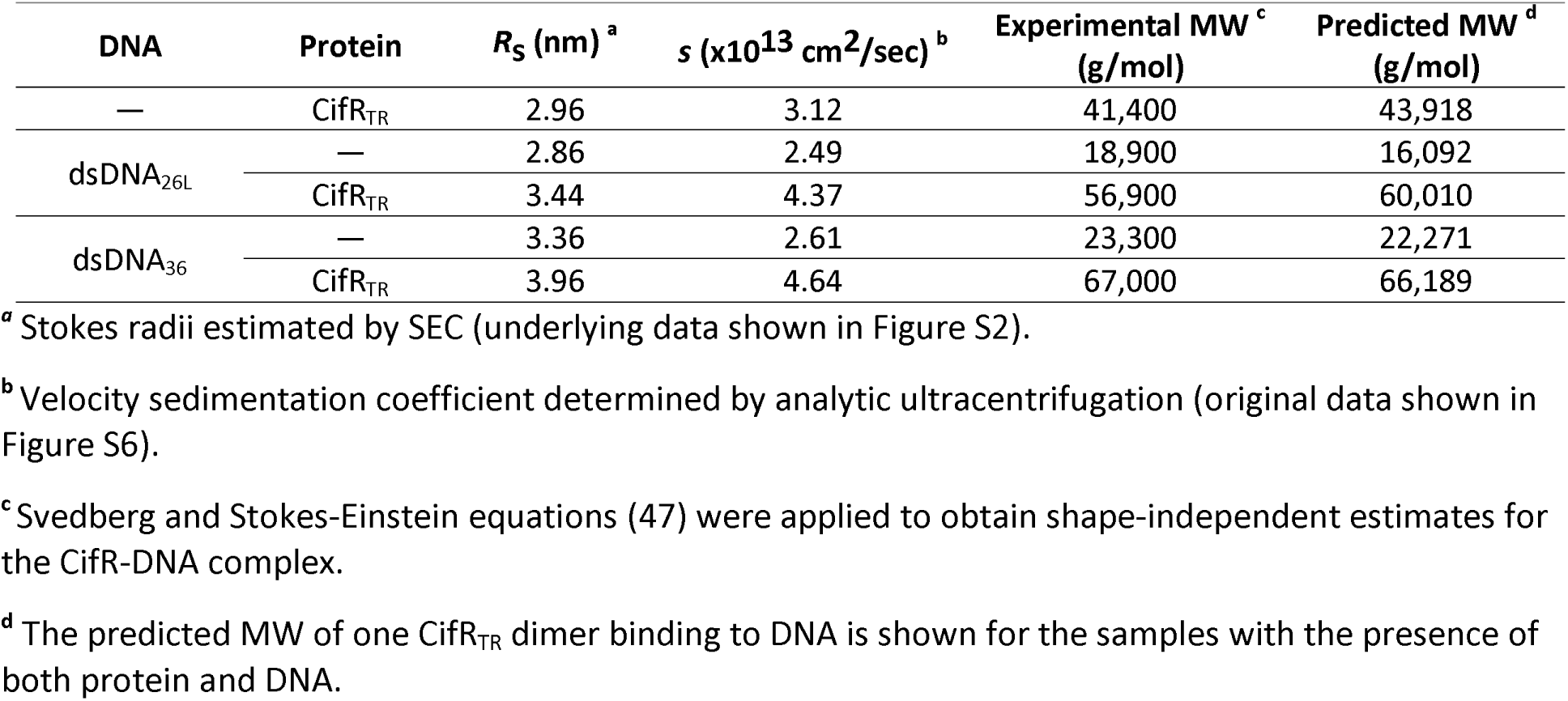
MW calculation of CifR_TR_-DNA complexes based on Stokes radius (*R*_s_) and sedimentation coefficient (*s*).

### The CifR_TR_:dsDNA_26L_ complex

To take advantage of the higher (2.6Å) resolution of the native dataset (Table 1), we used the CifR_TR_:dsDNA_26L_ complex structure from SeMet crystals as a search model for MR, and were readily able to determine initial phase estimates. Consistent with the improved diffraction, the quality of the electron density of the CifR_TR_:dsDNA_26L_ complex from this dataset was better than that of the SeMet dataset. However, the electron density for the apo-CifR_TR_ molecule was much weaker (Figure S4B) and did not support independent model building or refinement of the apo structure. As a result, only the CifR_TR_:dsDNA_26L_ model was refined against the native dataset.

Consistent with our hydrodynamic studies, the overall structure of the CifR_TR_:operator DNA complex is composed of one CifR_TR_ dimer and one dsDNA_26L_ duplex (Figure 4A, PDB ID: 6NSN). In each CifR protomer, three helices (α1 – α3) constitute the DNA-binding domain (DBD, green in Figure 4A), whereas helices α4 – α9 constitute the ligand-binding domain (LBD, pale cyan). The B-form DNA is slightly bent. The distance between the two Pro45 residues in the HTH motifs is 37.9 Å, allowing the motifs to sit in adjacent major grooves along one face of the DNA double helix. Meanwhile, the N-terminal extension that precedes the helix α1 in each CifR protomer inserts into the distal DNA minor groove on either side (Figure 4B).

**Figure 4.**
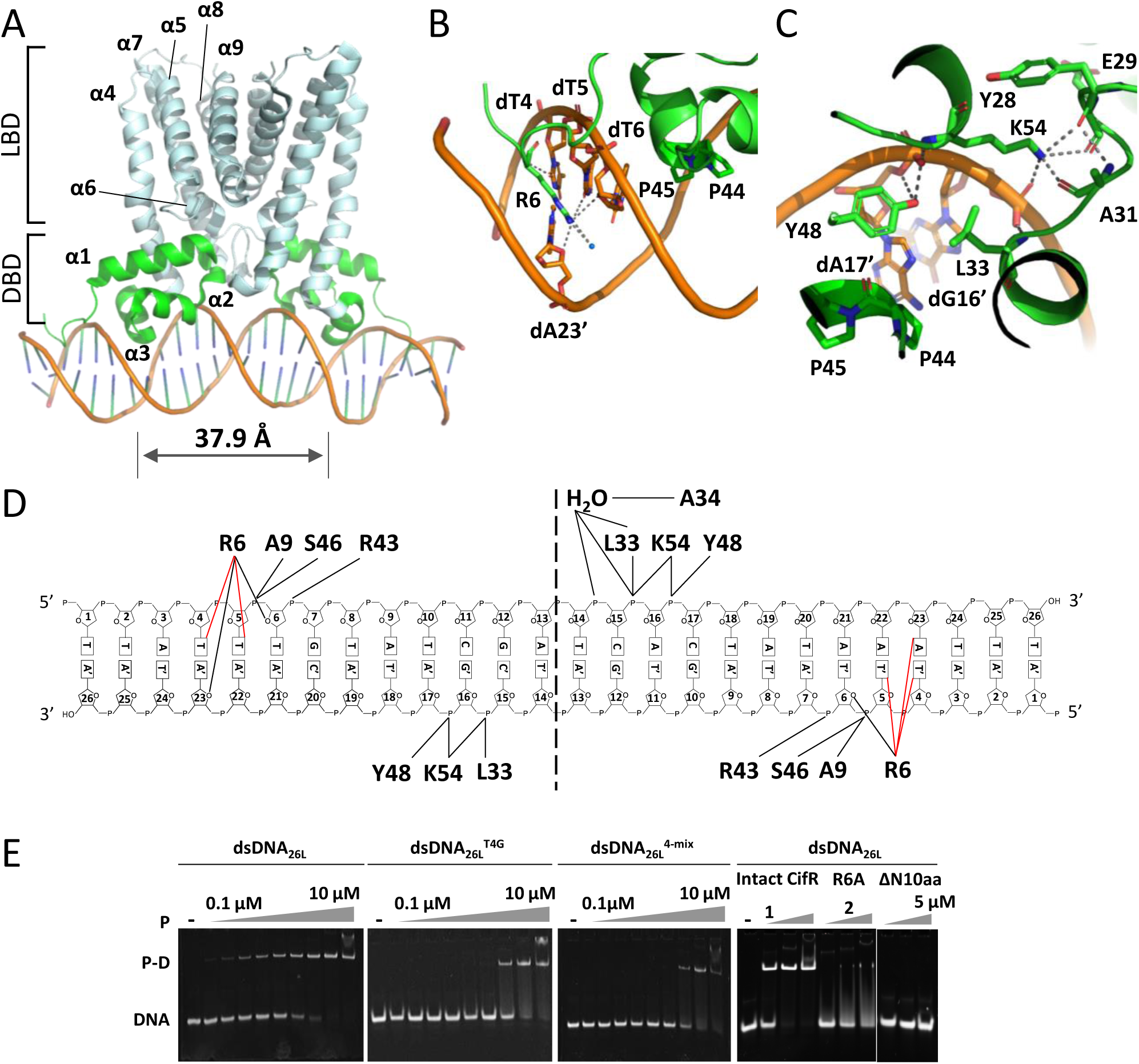
Structural analysis of the CifR_TR_-dsDNA_26L_ complex. **(A)** Overall structure of the CifR_TR_:dsDNA_26L_ complex. CifR_TR_ contacts the DNA duplex (orange) through the DNA-binding domain (DBD, green ribbon), while the ligand-binding domain (LBD, pale cyan ribbon) is distal to the protein-DNA interface. The nine helices that make up the bulk of the CifR_TR_ structure are labeled for one protomer. The distance between the two Pro45 residues in CifR dimer is shown. **(B)** Protein:DNA contact interface at the minor groove. Arg6 (green stick model) inserts into the DNA minor groove and forms hydrogen bonds (grey dashes) with several DNA bases (dT4, dT5, dT6, dA23’, orange sticks) and a water molecule (blue sphere). Pro44 and Pro45 are shown as green sticks. **(C)** Protein:DNA contact interface at the major groove. Leu33, Tyr48, Lys54 (green sticks) form hydrogen bonds (grey dashes) with backbone atoms of dG16’, dA17’ (orange sticks). Pro44 and Pro45 (green sticks) insert into the major groove but do not form hydrogen bonds. **(D)** Two-dimensional schematic of CifR-DNA interactions. The central dotted black line indicates the two-fold symmetry axis of the CifR dimer. Lines indicate interactions with DNA bases (red) or with the DNA backbone or a water molecule (black). **(E)** Functional analysis of protein-DNA contacts by EMSA. The DNA concentration was 0.5 µM. Protein concentrations tested were 0, 0.1, 0.2, 0.4, 0.8, 1, 1.6, 2, 4 and 10 µM for the nine titrations (panels 1-3), and 1, 2, and 5 µM for the three titrations (panel 4). Duplex dsDNA_26L_^T4G^ has mutations at T4 and A23. Duplex dsDNA_26L_^4-mix^ has mutations at C15T, A16C, C17T and T18G (see Table S1). R6A: CifR_TR_^R6A^; ΔN10aa: CifR_TR_ lacking residues 1-10.

Surprisingly, although operator specificity is usually encoded via base contacts in the major groove, CifR side chains only form contacts with the sequence-independent atoms of the DNA backbone (Figures 4C and 4D). Residues Leu33 from helix α2 and Tyr48 from helix α3 form hydrogen bonds with the phosphate groups of DNA bases A17’ and G16’, respectively, as does α4 helix residue Lys54 (Figure 4D), which is conserved among many TFR sequences (Figure S1). Lys54 also forms a hydrogen bond network with three residues (Tyr28, Glu29 and Ala31) located at the loop between helices α1 and α2. In contrast, there are no polar contacts between the HTH motif and the DNA bases in the major groove, which is typically the site that determines sequence specificity for TFRs (48). Water molecules can also bridge contacts between protein residues and the major groove, as seen for example, in the ms6564:DNA complex (49). However, in the CifR complex, the two residues most deeply inserted into the major groove are Pro44 and Pro45 (Figure 4B and 5C), neither of whose side chains can form hydrogen bonds. In fact, the only water-mediated hydrogen bond is formed by Ala34 from the α2 helix, but it also contacts only the phosphate groups of C15 and A16 (Figure 4D).

**Figure 5.**
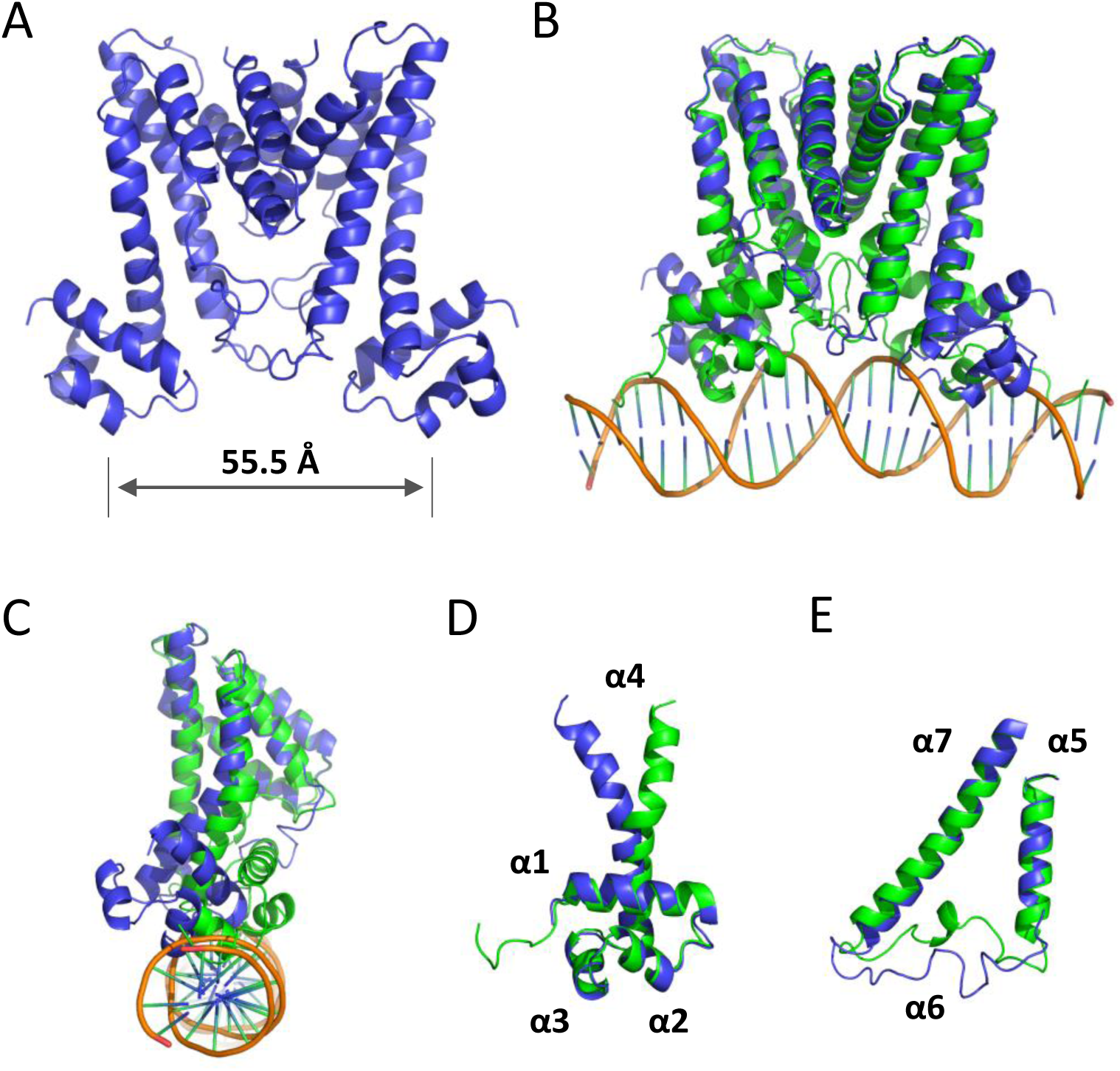
Comparison of DNA-bound CifR_TR_ and apo-CifR_TR_ structures. **(A)** Overall structure of apo-CifR_TR_. The structure of apo-CifR_TR_ is shown as blue ribbons. The distance between the two Pro45 residues in the CifR dimer is shown. **(B)** Superposition of the CifR_TR_:dsDNA_26L_ complex and the apo-CifR_TR_ dimer. C_α_/no outlier least-squares superposition of apo-CifR_TR_ (blue ribbons) and DNA-bound CifR_TR_ (green ribbons; orange DNA) performed in PyMol results in an RMSD of 3.1 Å. **(C)** Superposition of CifR_TR_:dsDNA_26L_ complex and apo-CifR_TR_ aligned at the ligand-binding domain of one protomer. A 90° rotation of the view in (B) results in the view shown here. Only one protomer is shown for clarity. **(D)** Conformational change of helix α4. Alignment of apo-CifR_TR_ and DNA-bound CifR_TR_ using the DNA-binding domain of one protomer of each dimer reveals the conformational change of helix α4 upon DNA binding. Only helices α1-α4 are shown for clarity. **(E)** Conformational change of helix α6. Alignment of the apo-CifR_TR_ and DNA-bound CifR_TR_ using the ligand-binding domain of one protomer of each dimer reveals the conformational change of helix α6 upon DNA binding. Only helices α5-α7 are shown for clarity.

Although there is no direct interaction between side chains of the HTH motif and any of the DNA bases, the binding affinity to CifR_TR_ was much lower when we mutated the four bases at the DNA major groove (C15T/A16C/C17T/T18G) (Figure 4E, third panel compared to the first). These central sequences may permit the DNA to form a cavity with a specific shape in the major groove to adopt the HTH motif of CifR_TR_. Also, we observed a noticeable degree of bending and kinking along the DNA where CifR binds (Figure 4A). It could be that this altered conformation of DNA seen with CifR_TR_ binding is facilitated by certain DNA sequences at the central DNA region.

### Minor groove contacts provide additional sequence specificity

In contrast to interactions involving the major groove, we observed significant protein-DNA base interactions in the minor grooves (Figure 4B and 4D). Structurally, the two external DNA minor grooves are rich in AT sequences. They are thus narrower by at least 2 Å than the minor grooves in the center of the dsDNA_26L_. Nevertheless, on both ends of the operator site, they accommodate the Arg6 side chain, which interacts with multiple bases (T4, T5, T6 and A23’). We tested the importance of these interactions by mutating the T4-A23’ and T4’-A23 to GC base pairs (Table S1) and we found that these substitutions led to significant decrease of complex stability (Figure 4E, second panel compared to first).

In a reciprocal experiment, the CifR_TR_-R6A mutant lost the majority of its DNA binding capacity (Figure 4E, fourth panel), despite exhibiting similar biochemical behavior to CifR_TR_, e.g., during purification. Very weak binding could be detected, and only at the highest protein concentration tested (5 μM). Deletion of the entire N-terminal loop of 10 aa completely abolished DNA binding (Figure 4E, fourth panel). This may reflect additional interactions between bp 1-3 and the very N-terminal residues. For instance, in chain A, Arg4 reaches DNA minor groove as well and likely interacts with T2, based on the limited electron density. We calculated the surface electrostatic potential using the Adaptive Poisson-Boltzmann Solver (APBS) (50) and found the expected positive values along the DNA-binding surface. The N-terminal loop, which interacts with the DNA minor groove, exhibits very strong positive potential as well (Figure S7A), primarily due to the presence of two arginine residues (Arg4 and Arg6). These results together suggest that minor groove contacts provide critical sequence specificity for CifR_TR_-operator recognition.

### Structural comparison of CifR_TR_ in the presence and absence of DNA

The overall structure of apo-CifR_TR_ (Figure 5A, PDB ID: 6NSR) reflects a distinct conformation compared to that seen in complex with DNA (Figure 4A, PDB ID: 6NSN). A C_α_ least-squares superposition of the apo-CifR_TR_ dimer and DNA-bound CifR_TR_ dimer yields an RMSD of 3.1 Å (Figure 5B). The RMSD of the individual DNA-binding domains (helices α1 – α3) from each monomer is 0.75 Å (Figure 5D), while that of each ligand-binding domain (helices α5 – α9) is 0.71 Å (Figure 5C). Thus, the structure of each individual domain is conserved. However, their relative orientation has changed. The RMSD for the α4 helix, which connects the two domains, is 2.0 Å, primarily reflecting its enhanced curvature in the apo structure (Figure 5D). As a result, the two HTH motifs are further apart from the dimer axis in the absence of the DNA (Figure 5C): the distance between the Pro45 residues in each of the two HTH (55.5 Å; Figure 5A), has increased 17.6 Å relative to the CifR_TR_:dsDNA_26L_ complex (Figure 4A).

While the ligand-binding domain is relatively conserved, the conformational changes observed for the α6 helix (residues 100-120) were far higher than for any other secondary structural element, with an RMSD of 5.0 Å (Figure 5E). In part, this reflects a significant displacement of the dominant conformation. In part, it also reflects the weakness of the corresponding electron density (Figure S8D) in the apo-protein structure, most likely reflecting conformational flexibility in the absence of bound operator DNA, compared to other helices (e.g., the α6 helix; Figure S8B) and the corresponding helices in the DNA-bound structure (Figures S8A and S8C).

Taken together, the conformational changes associated with DNA binding also have dramatic effects on the overall compactness of the protein fold. In the DNA-bound state, cavity simulation using CASTp (51) revealed two potential ligand-binding pockets of CifR_TR_ (pockets 3 and 4 in Figure S9A) surrounded by helices α5-α7 in each protomer. These pockets have high positive surface potential (Figure S7B, inset). Two additional pockets were detected beneath α6 (pocket 1 and 2, Figure S9A), which may facilitate effector entry into the binding pocket. If so, the differential flexibility of the α6 helix could be coupled to ligand entry and exit. In the apo state, the distance between the protomer HTH motifs increases by nearly 18 Å (Figure 5A) relative to the DNA-bound state (Figure 4A). This creates a relatively large volume for ligand entry between the domains in which the four isolated pockets are connected to each other (Figure S9B).

### The role of Cys107 in ligand recognition

As we observed from the TFR sequence alignment above (Figure S1), the DNA-binding domain is relatively conserved among TFRs (Figure S1). We mapped the conserved residues to the CifR_TR_:dsDNA_26L_ complex structure and found that most of these residues are located at the interface formed among helices α1 – α3 and the N-terminus of helix α4 (Figure S10A).

In contrast, the C-terminal ligand-binding domain shows very little conservation throughout (Figure S1, Figure S10A). This polymorphism may correspond to the wide variety of ligands that are recognized by TFRs. However, there is a small group of residues in the ligand-binding domain that are conserved in over 40% of sequences. These include Lys54, Leu57, Gly106, Cys107, Gly149, Ala161, and Gly170. The residues fall into three groups: the α4 helical residues, the C-terminal residues, and the ligand-binding residues.

First, the conserved residues Lys54 and Leu57 are located in helix α4 which forms a connection between the DNA-binding domain (helices α1 – α3) and the ligand-binding domain (helices α5 – α7) for most TFRs (52). As discussed above, in the structure of CifR_TR_:dsDNA_26L_ complex, Lys54 forms a polar contact with the DNA backbone as well as with three residues (Tyr28, Glu29 and Ala31) located at the loop between the α1 and α2 helices (Figure 4C). Leu57 forms hydrophobic interactions with Leu36 and Leu47 from α2 and α3, respectively (Figure S10B). Thus, these two residues probably have a role in maintaining the relative positions of the DNA-binding domain and helix α4.

Second, the most C-terminal conserved residues (Gly149, Ala161 and Gly170) are either located in helix α8 or in the loop between helices α7 and α8. The α8 helix constitutes the protein dimerization interface (Figure 5B). Therefore, these three conserved residues may well be important for protein dimerization.

The third group consisted of the remaining two conserved residues, Gly106 and Cys107. These residues are adjacent to the α6 helix, which is part of the ligand-binding triangle of most TFRs (52), composed of helices α5 – α7, as shown in Figure 6A and 6B. Gly106 takes on a glycine-specific main-chain torsional configuration, which may be necessary to achieve the observed side-chain orientation of Cys107. Given its position at the center of the LBD and the interesting chemistry of sulfhydryl groups, we hypothesized that Cys107 might play a role in ligand recognition. We tested this by replacing Cys107 with either a serine or a threonine, or by treating CifR_TR_ with a cysteine-modifying compound.

**Figure 6.**
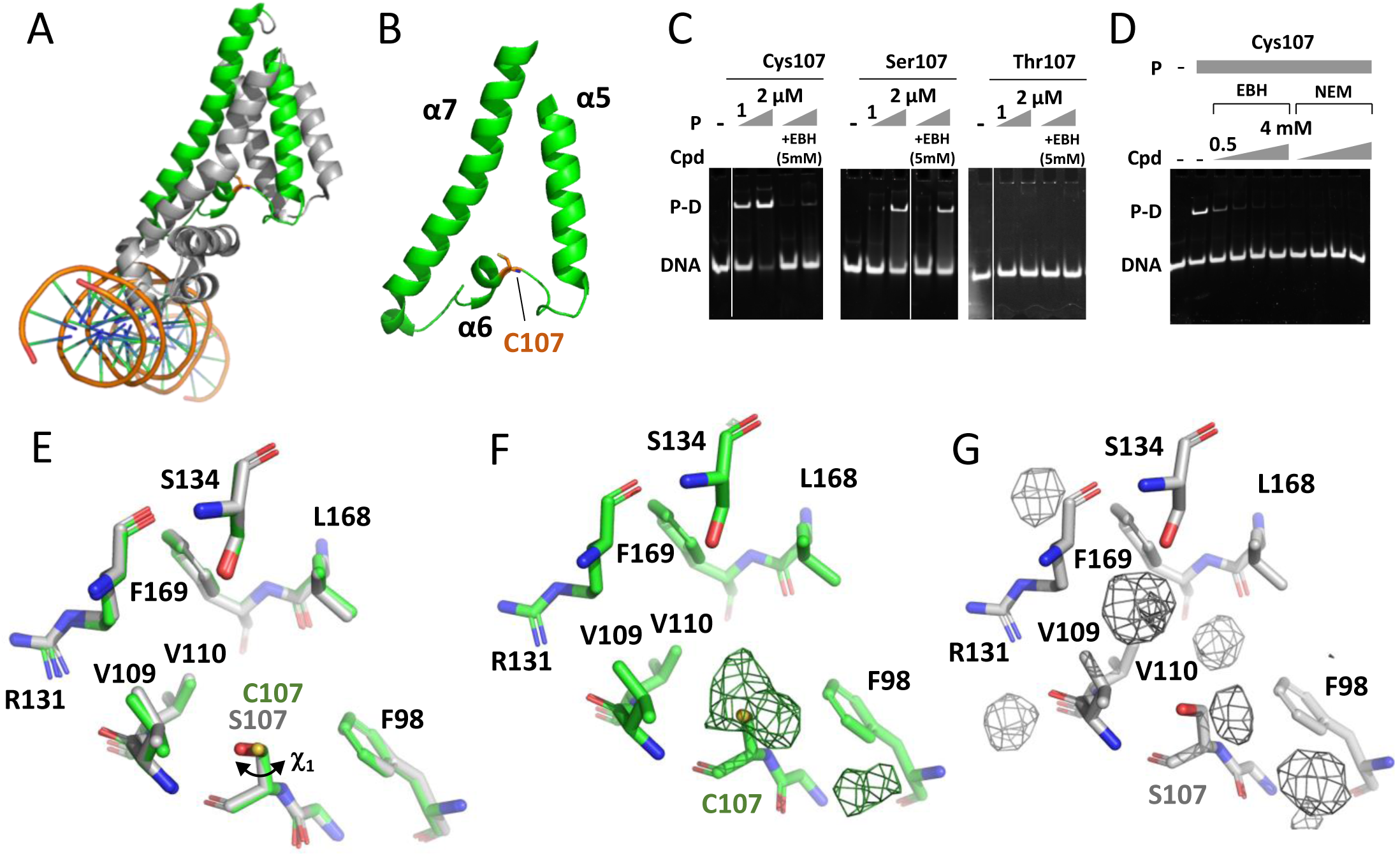
The role of conserved residue Cys107 in the potential epoxide-binding pocket. **(A)** The potential epoxide-binding pocket in CifR_TR_. Only one CifR_TR_ protomer is shown in the CifR_TR_:dsDNA_26L_ structure for clearer view of the potential epoxide-binding pocket formed by helices α5 - α7 (green); other helices are shown in grey. **(B)** A closeup view of the proposed epoxide-binding pocket. Cys107 lies at the bottom of the epoxide-binding pocket (orange stick figure). **(C)** Response of CifR_TR_ Cys107 mutants to EBH. The involvement of Cys107 in ligand recognition and/or transduction of ligand binding to DNA release was assessed by EMSA with CifR_TR_, CifR_TR_^C107S^, and CifR_TR_^C107T^ mutants. The DNA concentration was 0.5 µM. Cpd = compound. Solvent (DMSO) alone did not disrupt protein-DNA complexes (not shown). **(D)** Response of CifR_TR_ protein to NEM analyzed by EMSA. A mixture containing 0.5 µM DNA and 1 µM CifR_TR_ was titrated with 0.5, 1, 2 and 4 mM EBH or the cysteine-modifying compound NEM. **(E)** Comparison of the ligand-binding pockets of CifR_TR_:dsDNA_26L_ and CifR_TR_^C107S^:dsDNA_26L_. DNA-bound CifR_TR_ (green sticks) and DNA-bound CifR_TR_^C107S^ (light grey sticks) were aligned using the ligand-binding domains. **(F)** and **(G)** m*F*_O_-D*F*_C_ maps of the ligand-binding pockets of DNA-bound CifR_TR_ (green sticks) and DNA-bound CifR_TR_^C107S^ (grey sticks). Positive peaks are shown as green mesh for CifR_TR_:dsDNA_26L_ and grey mesh for CifR_TR_^C107S^:dsDNA_26L_. Both maps are contoured to 3.0σ.

CifR_TR_^C107S^ contains a sterically similar but chemically distinct side chain. While CifR_TR_^C107S^ displays a reduced binding affinity to operator DNA as compared to CifR_TR_ (Figure 6C), the residual binding exhibited by the C107S mutant is also resistant to the addition of epoxide EBH (Figure 6C), implicating residue Cys107 in epoxide responsiveness.

To ensure that the mutation did not fundamentally disrupt the structural integrity of the CifR transcription factor, we crystallized the CifR_TR_^C107S^:dsDNA_26L_ complex and determined its structure at a resolution of 2.8 Å (PDB ID: 6NSM, Table 1). The structure is very similar to that of the CifR_TR_:dsDNA_26L_ complex, with an RMSD (0.15 Å) comparable to the estimated coordinate error of the structure itself (0.21Å). The side chains pointing into the ligand-binding pocket of these two structures are shown in Figure 6E. The Ser107 hydroxyl group has reoriented compared to the Cys107 thiol group; χ_1_ shifts by approximately 35° (Figure 6E). It is worth noting that in the CifR_TR_:dsDNA_26L_ complex structure, we observed some positive density in the difference map near the thiol group of Cys107 (Figure 6F). Due to the resolution, we could not determine the source of this electron density, but it might represent one or more covalent modifications of Cys107. Some difference electron density peaks are similarly observed in the vicinity of residue 107 in the CifR_TR_^C107S^:dsDNA_26L_ structure. However, these densities are not contiguous with the electron density for the replacement serine residue (Figure 6G). The pattern of difference density peaks is thus compatible with covalent and non-covalent ligand binding, consistent with the differential reactivities of thiol and hydroxyl groups, respectively.

Both the mutant CifR_TR_^C107T^ and the chemically modified cysteine in CifR_TR_ displayed similar properties to each other. CifR_TR_^C107T^ has lost its DNA binding ability completely (Figure 6C). The difference between serine and threonine is the addition of one γ-methylene group, which may mimic the ligand-bound state. In addition to genetically editing residue Cys107, we applied the cysteine-modifying compound NEM to the CifR_TR_:dsDNA_26L_ complex. Even at the lowest concentration tested (0.5 mM), it completely disrupted the DNA-binding capacity of CifR_TR_ (Figure 6D). The covalent modification of Cys107 might structurally mimic the ligand-bound state, which leads to a conformational change and release from the DNA. Taken together, these results indicate that the conserved Gly106-Cys107 pair is likely involved in ligand recognition and the transduction of ligand-binding to DNA release for this novel epoxide-sensitive regulator.

## DISCUSSION

### CifR:operator interactions

TetR family transcriptional regulators (TFRs) are found across a wide range of bacterial species and in response to different ligands regulate diverse physiological functions, including drug resistance, metabolism, and signaling processes (48). Compared to the large number of known family members, only a relatively small number of TFRs have been characterized mechanistically (53–57). As outlined below, the structures of CifR alone and in complex with DNA share some fundamental features of the TFR family, but also illuminate some novel characteristics. Collectively, the structures of TFR:DNA complexes reveal two fundamentally distinct stoichiometries, with a single operator duplex binding either one or two TFR dimers. Sites as short as 14-15 bp appear to represent the minimum footprint that can comfortably accommodate a single TFR dimer (Figure 7A) (41), but can also accommodate two TFR dimers (e.g., SlmA; Figures 7C and S3) (42, 48). Longer operator sequences are observed, and can harbor either one (e.g., DesT; Figures 7C and S3) or two (e.g., QacR; Figures 7B and S3) dimer-binding sites.

**Figure 7.**
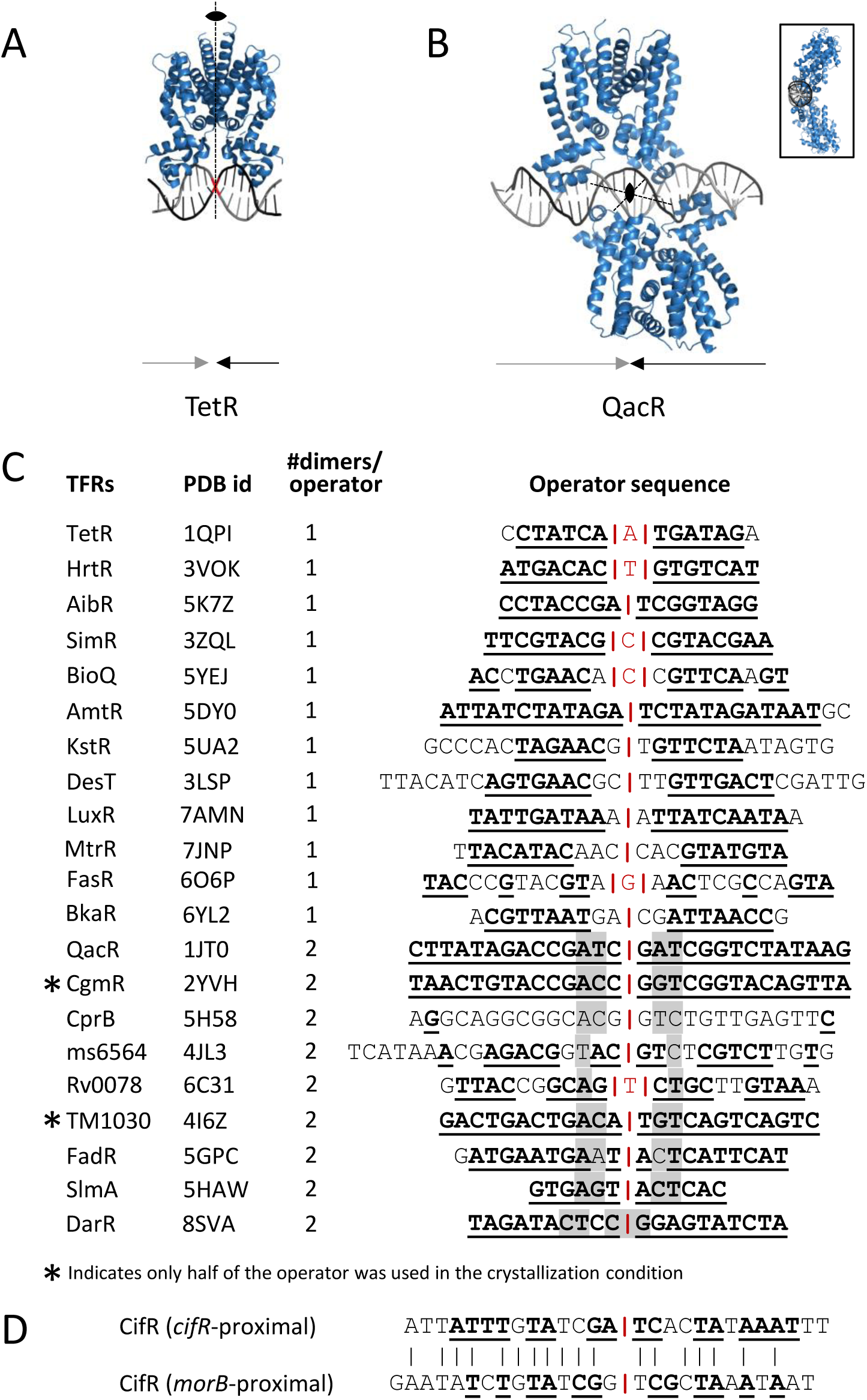
Recognition of operator DNAs by TFRs. **(A)** and **(B)** Proteins are shown as blue ribbons. DNA is shown in black and grey for each strand of the DNA duplex. The solid ovals indicate the two-fold rotational symmetry with the dotted lines indicating the orientation of the axes. The arrows below indicate the inverted repeats of DNA. **(A)** Structure of DNA-bound TetR. The center of the DNA is shown in red. **(B)** Structure of DNA-bound QacR. The rotation axis is perpendicular to the plane of view. The inset shows a 90°-rotated view of the main panel. **(C)** Operator DNA sequences of 21 TFRs with known DNA-complexed structure. The operator DNA sequences analyzed here are those used in the protein-DNA complex structure determination. The predicted center of each inverted repeat is shown with red text or bars. Structures in which only half of the operator DNA was used in the crystallization are labeled with an asterisk. The bases with potential inverted repeats property are underlined and in bold. The centers for two embedded repeats are highlighted in grey. The embedded repeats are shown in detail in Fig. S11. **(D)** Operator DNA sequences of CifR. The bases with potential inverted repeats property are underlined and in bold. The conserved residues between the two operator DNA sequence are marked by vertical alignment bars (black).

The operator DNA sequences of most TFRs form inverted repeats (Figure 7C). Among known structures with two dimers bound to an operator, the axis of each dimer almost always aligns not with the center of the overall inverted repeat, but rather with the centers of less symmetric embedded repeats located to either side. In either case, sequence repeats generally reflect the 2-fold rotational symmetry of the TFR dimer (58). The centers of the dual embedded repeats are highlighted with grey shadows in Figure 7C and the embedded repeats are shown in detail in Figure S11. The presence of embedded repeats in the operator DNA thus can serve as a clue for a two-dimer binding mode. Indeed, mutagenesis to create an embedded repeat has been leveraged to engineer enhanced affinity of protein:operator interactions in the DarR system (59).

Compared to most other TFR operator DNAs (Figure 7C), the *cifR*-proximal and *morB*-proximal sites do not show strong embedded-repeat signatures (Figure 7D). They are relatively long and show considerable sequence divergence from each other (Figure 7D). However, not all dual-site operators show clear embedded repeat signatures (e.g., CprB and DarR; Figures 7C and S11), and in the case of DarR binding to the wild-type operator, one of the dimers actually aligns with the center of the main inverted repeat, while the other is offset by 3 bp (59). Thus, it was difficult to predict based on sequence alone whether one or two CifR dimers would form a complex with the operator. However, our structural and biochemical data clearly show that apo-CifR_TR_ is a dimer and that the CifR_TR_:DNA complex is composed of one dimer bound to one dsDNA duplex.

Since these structures were first determined, there have been significant advances in the computational modeling of protein complexes (60–62). Since the structures reported here have been publicly available for training, CifR is not a strong test case for the ability to predict TFR conformations or their interactions with DNA. However, we were surprised to note that the AlphaFold Protein Structure Database (63) still predicts a monomeric structure for the PA14 CifR sequence, despite having identified homology to the deposited PDB structure. We confirmed that the current version of AlphaFold can successfully predict a dimeric structure for the CifR sequence (Figure S12), although in the absence of DNA it predicts a conformation closest to the DNA-bound conformation, differing primarily at the N-terminal recognition motif. Indeed, it is widely recognized that the accurate computational modeling of protein:DNA complexes remains challenging for a number of reasons. Nonetheless, recent advances raise the prospect of more accurate predictions in the near future (64). For these studies, paired structures of DNA-binding proteins alone and in complex with physiologically relevant operator sites, as seen here for CifR, will provide important opportunities for training and validation.

### Lack of major-groove hydrogen-bond interactions

TFR HTH motifs are usually positively charged, which facilitates operator binding given the negatively charged phosphate backbone of DNA. The expected positive surface potential was observed along the DNA-binding surface of CifR_TR_ with enhanced positive charge at the N-terminal loop, which interacts with the DNA minor groove (Figure S7A). In addition to general electrostatic attraction, the specific recognition of operator sequences usually involves the formation of hydrogen bonds between DNA bases and protein residues. The HTH motifs of TFRs typically insert into the DNA major groove, where base-specific hydrogen-bonding partners are more fully exposed for stereochemical recognition. Consistent with the canonical view, mutations of the bases in the DNA major groove near the HTH motif led to significant reductions in the binding affinity of CifR_TR_ (Figure 4E, fourth gel). This indicates that the major-groove sequence plays a role in CifR_TR_ recognition.

However, compared to the operator-bound structures of other TFRs, the CifR:DNA complex is distinguished by a lack of hydrogen bonds between the HTH motif residues and DNA bases in the major groove. One reason is that the two residues most deeply inserted into the major groove are prolines, which are hydrophobic residues. This may explain why the two operator DNA sequences of CifR exhibit only very weak inverted repeats (Figure 7D), as they are not strongly constrained by individual stereochemical contacts.

One potential source of additional stereochemical specificity involves hydrophobic interactions between the two proline residues and adjacent DNA bases. Another possibility is that the DNA sequence forms a specific cavity or permits DNA bending to accommodate the HTH motif of CifR_TR_. The dsDNA_26L_ duplex in the complex was slightly bent compared with standard B-form DNA. Such deformation is typically sequence-dependent, and if required for transcription-factor binding, represents a form of “indirect readout” (65–67).

### Compensatory minor-groove interactions

In addition to evidently indirect recognition in the DNA major groove, we found that the N-terminal extension inserts into the DNA minor groove, where Arg6 interacts directly with multiple bases (Figure 4A-4C), where the minor groove narrows noticeably. Rohs and colleagues have reported that a narrowed minor groove is often associated with AT-rich sequences, like those found in dsDNA_26L_, and that such narrowing is frequently associated with binding of arginine side chains (68). Altogether, the combination of the hydrophobic or shape-recognition mechanism at the major groove and the direct base-sequence read-out at the minor groove appear to constitute the complete recognition mechanism between CifR_TR_ and its operator DNA.

In fact, CifR is not the only TFR:DNA complex that exhibits specific base interactions at the minor groove. Six previously determined TFR:DNA structures show a comparable interaction in the DNA minor groove as well. This list includes the global nitrogen regulator AmtR (69), a master regulator ms6564 (49), the efflux pump regulators SimR (44) and EilR (70), the quorum-sensing regulators CprB (43) and LuxR (54). For AmtR, SimR, CprB, and LuxR, arginine residues are also found to interact with the bases in the minor groove (Figure S13A-D). Contacts are mediated by an arginine and a serine in ms6564 and by a tyrosine residue for EilR (Figure S13E and F).

We believe that many TFRs may exploit this recognition strategy. The sequence alignment of 938 TFRs (Figure S1) shows that Arg6 is relatively highly conserved. Also, Le *et al*. have analyzed the sequences of 12,715 non-redundant TFRs and found that 72% of them have an N-terminal extension longer than 11 amino acids (44). Furthermore, residues upstream of the HTH motif are often not resolved in TFR complex structures, including several where a longer operator DNA binds a single homodimer, leaving the minor groove exposed at either end. BioQ was co-crystallized in complex with a 19-bp operator, and the footprint of the TFR covers the length of the duplex, which presumably represents the extent of a canonical TetR binding site. In comparison, KstR was crystallized as a single dimer in complex with a 26-bp duplex. Although the 5’ and 3’ base pairs did not appear to form direct protein contacts, the first 30 aa of KstR could not be located in the experimental electron density. The DesT:operator complex was co-crystallized using a 31-bp construct. However, several DNA bp were not resolved on either end of the duplex, and three N-terminal side chains are missing in the structure. These results together suggest that the interaction between N-terminal extension and DNA minor groove could be a relatively common type of interaction among TFRs. Because electrostatic interactions are not directional, in some cases, N-terminal residues may be able to contribute to affinity without requiring a defined binding poise.

### Co-crystallization of CifR-DNA complex and apo-CifR

A very unusual feature of our structural results is that we were able to capture a DNA-binding protein in two different states within a single crystal lattice. CifR_TR_ retained some tendency to precipitate in the absence of DNA but retained much higher solubility and stability in the presence of DNA, like many DNA binding proteins. Fundamentally, crystallization requires regular lattice packing, which usually requires samples to be homogenous. In fact, in an attempt to drive the thermodynamic binding equilibrium towards complex formation, we mixed the CifR_TR_ protein with a 1.25-fold molar excess of dsDNA_26L_ compared to the dimer protein. We attempted to obtain the structure of the complex as our initial goal. We were therefore particularly surprised to discover a second dimer, apo-CifR_TR_, caged in the bulk solvent channel of a lattice determined by CifR_TR_-DNA complex. Residues Tyr48 and Lys54, which are involved in DNA-binding, are therefore free in apo-CifR_TR_, and they are located at the interface with the adjacent DNA-bound CifR_TR_ molecule (Figure 3B, bottom left). Thus, our hypothesis is that the lattice-binding site was able to compete with DNA for the binding of apo-CifR_TR_. DNA binding may also have been disfavored by the crystallization buffer, that required the presence of 200 mM magnesium chloride to form well diffracting crystals. We have no explanation for why the occupancy of apo-CifR_TR_ is higher in the crystals formed by the SeMet labeled protein. No methionine side chains directly participate in CifR crystal lattice formation, although SeMet labeling is known to affect protein solubility and crystal stability (71, 72). In any case, the dual-structure lattice provided us with the unexpected opportunity to visualize the conformational effects of interactions with the operator site.

### CifR conformational changes associated with DNA binding

There are both local and global structural differences between the apo and DNA-bound forms of CifR. Specific changes are observed in helices α4 and α6 of the ligand-binding domain, which are otherwise well conserved. The α4 helix connects the ligand-binding domain and HTH motif. In both structures, α4 is bent in the middle, but the angles of curvature are noticeably different. The position of the C-terminal half of α4 relative to the ligand-binding domain is conserved between the apo and the DNA-bound states (Figure 5B), as is the position of the N-terminal half relative to the DNA-binding domain (Figure 5D). This flexing at the middle of the α4 helix is the key to the conformational rearrangement between the domains. Our hypothesis is that bending of the α4 helix mediates conformational crosstalk between the effector and DNA binding sites that ultimately controls Cif operon transcription.

Among the nine helices in apo-CifR_TR_, α6 exhibits the poorest electron density (Figure S8D), which suggests flexibility in the absence of ligand or DNA. As shown above, apo-CifR_TR_ can form dimers in the absence of reducing agent (Figure 1E). It is likely that the flexibility of the α6 enables the embedded Cys107 side chains of each monomer to interact covalently. Within the PDB, the structure with the closest sequence identity to CifR is ID 2i10, a putative TFR. Even though the structure was determined at a resolution of 2.0 Å, part of the α6 helix is missing in the refined structure, whereas the other helices are all very well defined experimentally. In contrast, in the DNA-bound structure of CifR_TR_, α6 is well defined (Figure S8C).

Conformational changes involving both helices have also been observed in other TFR structures. In the case of TetR, the α6 helix opens its C-terminal turn upon ligand binding, which in turn shifts the position of the α4 helix (73). For QacR, ligand binding leads to a movement of the α6 helix, which causes a pendulum motion of the α4 helix (74). In both cases, these changes are communicated to the DNA-binding domains. However, this is not the unique mechanism for allosteric regulation. For CprB, effector binding reorganizes the dimerization interface, which in turn alters the relative position of the two HTH motifs and leads to DNA release (43, 75). For CifR, the dimer interface remains well conserved (Figure 5B).

The conformational changes associated with DNA binding also have dramatic effects on the overall compactness of the protein fold. Complexing with its operator DNA compacts CifR_TR_ protein conformation (Figures S8A and S8B). Even accounting for the 60% occupancy of apo-CifR_TR_ within the lattice, the apo structure exhibits higher B-factors than does DNA-bound CifR_TR_ (Table 1). It is reasonable to hypothesize that DNA binding reduces conformational flexibility and thus allosterically competes with ligand binding to sites which are preferentially accessible in the apo state.

### Potential biomedical relevance

Epoxides play important roles in cell physiology. They can be generated during catabolic processes (76) and are also used as chemical signals between microbes (e.g., fosfomycin) (13, 77). In human cells, polyunsaturated fatty acids can be enzymatically epoxidated, yielding compounds that play diverse roles in immune system signaling (7). Our earlier work showed that microbes can regulate transcription of a virulence circuit in response to the presence of epoxides. In *P. aeruginosa*, CifR controls the expression of the *cif* operon, which encodes Cif, a secreted epoxide hydrolase that dysregulates host immune responses and facilitates infection. We are developing small-molecule and nanobody inhibitors of Cif EH activity (78–80). However, the availability of structural and biochemical information on CifR may provide opportunities to block the ability of *P. aeruginosa* to recognize epoxides, mimicking the behavior of the C107S effector-insensitive mutant. This would suppress epoxide-mediated expression not only of *cif*, but also of the two other members of the operon: a major facilitator symporter homolog (*mfs*) and a morphinone reductase (*morB*) (16). While their roles have not been characterized in *P. aeruginosa*, MFSs are very often involved in critical cellular physiological process of bacteria (81, 82). In addition, such studies will provide important insights into the mechanism by which epoxide binding triggers the release of operator DNA in CifR and will increase our molecular understanding of this therapeutically relevant family of bacterial transcription factors. Furthermore, prokaryotic transcription factors can be engineered to activate established cellular response pathways in response to new small-molecule signals (83, 84). Our characterization of the CifR epoxide-sensing circuit may expand this toolbox to include a new class of chemical scaffolds. Overall, the molecular structures of CifR helped us to understand its mechanism of operator recognition and uncovered critical residues for ligand sensing. These results offer new insights into the stereochemical regulation of an epoxide-based virulence circuit in critically important clinical pathogen.

## Data availability

For all structures reported here, atomic coordinates and structure factors have been deposited with the Protein Data Bank. See Table 1 for relevant identifiers.

## Acknowledgments

We would like to thank Drs. Jean Jakoncic, Alexei Soares and Dieter Schneider at Beamline 17-ID-1 of NSLS-II at BNL, Drs. Martin Fuchs and Wuxian Shi at Beamline 17-ID-2 of NSLS-II at BNL, Drs. Aina Cohen and Michael Soltis at Beamlines 9-2 and 12-2 of SSRL at SLAC, and Drs. Nagarajan Venugopalan and Michael Becker at Beamline 23-ID-D GA/CA of APS for help with data collection. We would like to thank Adam H. Barczewski for helping with the data collection on CifR_TR_^C107S^:dsDNA_26L_ complex. We would like to thank Christopher Pennil for preliminary experiments and helpful discussions and Adam Simard for assistance with the graphical abstract.

