## Supplementary figures for "Molecular basis for the transcriptional regulation of an epoxide-based virulence circuit in *Pseudomonas aeruginosa*"

### Figure S1

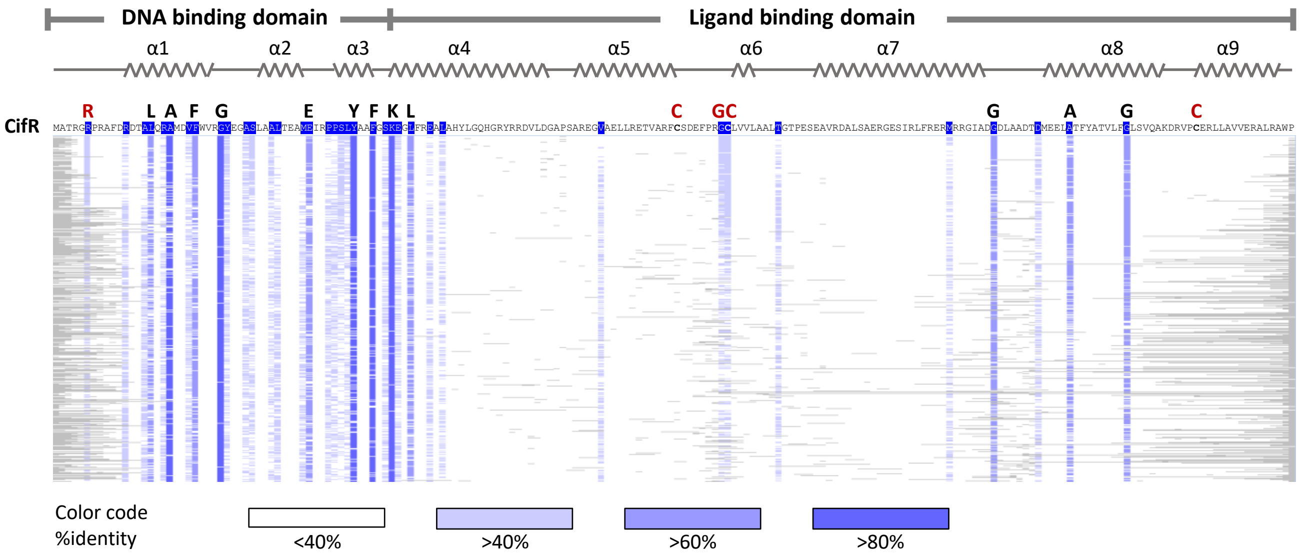


**Figure S1.** CifR protein domain organization and sequence conservation. The domain organization and experimental secondary structure derived from DSSP analysis of the crystal structure determined in this study are shown above the results of a PSI-BLAST (Position-Specific Iterative Basic Local Alignment Search Tool) analysis involving 938 TFRs with the highest sequence similarity to CifR from the NCBI protein database. The sequences vary from 100 to 196 amino acids in length, with grey horizontal lines indicating alignment gaps. The columns of varying shades of blue under each CifR residue indicate the percent conservation of that residue among the aligned TFRs. Residues with higher than 60% conservation (black) are shown in a larger font above the CifR sequence, together with other residues discussed in the text (red).

### Figure S2

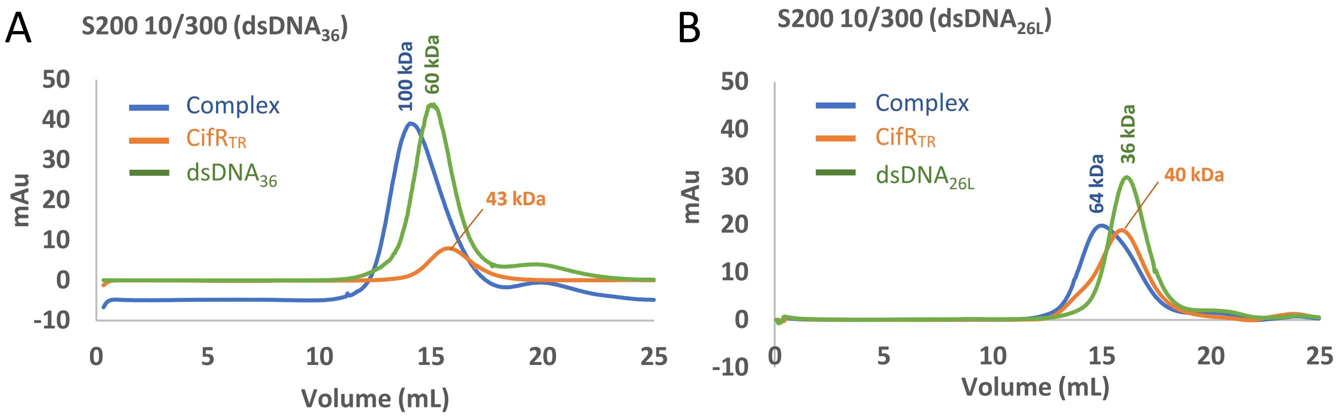


C

| Protein | $V_e$ (10/300) | MW | $R_S$ (nm) | $K_{av}$ (10/300)[a] | $(-\log K_{av})^{1/2}$ |
|---|---|---|---|---|---|
| **Blue dextran ($V_o$)** | 8.78 | | | | |
| **Thyroglobulin** | 10.05 | 670000 | 8.58 | 0.086 | 0.533 |
| **γ-globulin** | 12.78 | 158000 | 5.1 | 0.271 | 0.284 |
| **Ovalbumin** | 16.04 | 44000 | 2.8 | 0.491 | 0.154 |
| **Myoglobin** | 18.55 | 17000 | 1.9 | 0.661 | 0.090 |
| **Vitamin B12** | 22.18 | 1350 | | 0.907 | 0.021 |
| **$V_{total}$ ($V_t$)** | 23.56 | | | 1 | 0 |

[a] $K_{av}=(V_e-V_o)/(V_t-V_o)$

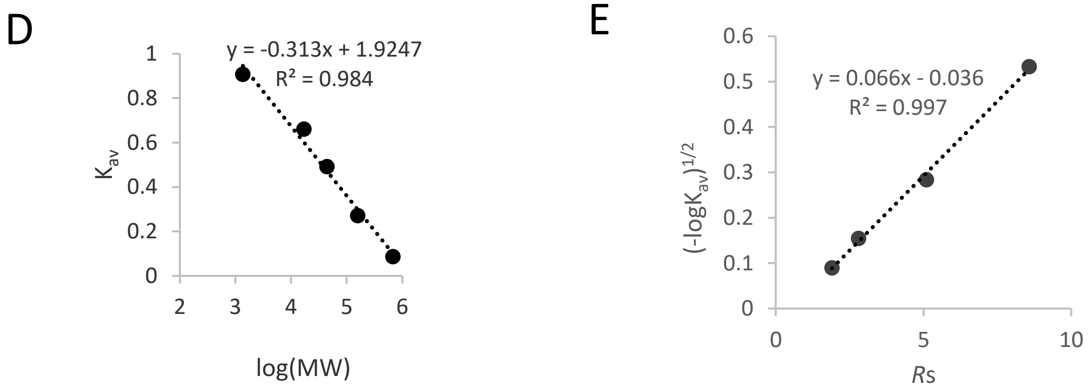


**Figure S2.** Determination of $CifR_{TR}$:DNA hydrodynamic behavior via size-exclusion chromatography. Size-exclusion chromatography elution profiles of protein, DNA, and protein-DNA complex using *cifR*-prox $dsDNA_{36}$ **(A)** and *cifR*-prox $dsDNA_{26L}$ **(B)**. The retention times of $CifR_{TR}$ are consistent with the predicted molecular mass of a protein dimer (40-43 kDa). The elution profile of the $CifR_{TR}$:$dsDNA_{26L}$ complex yielded an estimate (64 kDa) close to that expected for a single dimer bound to the operator. In contrast, the elution profile of the $CifR_{TR}$:$dsDNA_{36}$ complex yielded a significantly higher molecular mass estimate (100 kDa), with a value close to that expected for a complex formed by two CifR dimers bound to a single operator (109 kDa). The $dsDNA_{36}$ eluted at a volume corresponding to 60 kDa, compared to a calculated molecular mass of 23 kDa. The earlier retention time is likely due to the elongated nature of DNA, since SEC retention time depends on hydrodynamic radius, which is shape dependent. Calibration standards are globular proteins, and the extended nature of the DNA double helix can thus confound molecular mass determination (see also Table 2). **(C)** Characteristics of molecular-weight standards used to calibrate Superdex 200 10/300 column. **(D)** $K_{av}$ values of the standards were plotted against log(MW) and fit by linear regression (equation). This calibration was used to assign approximate molecular weights to each elution peak shown in (A) and (B). **(E)** A second standard curve was generated by linear regression fitting of $(-\log K_{av})^{1/2}$ values plotted against their known Stokes radii. The Stokes radii shown in Table 2 for DNA, $CifR_{TR}$, and $CifR_{TR}$:DNA complexes were calculated from the resulting equation.

Figure S3

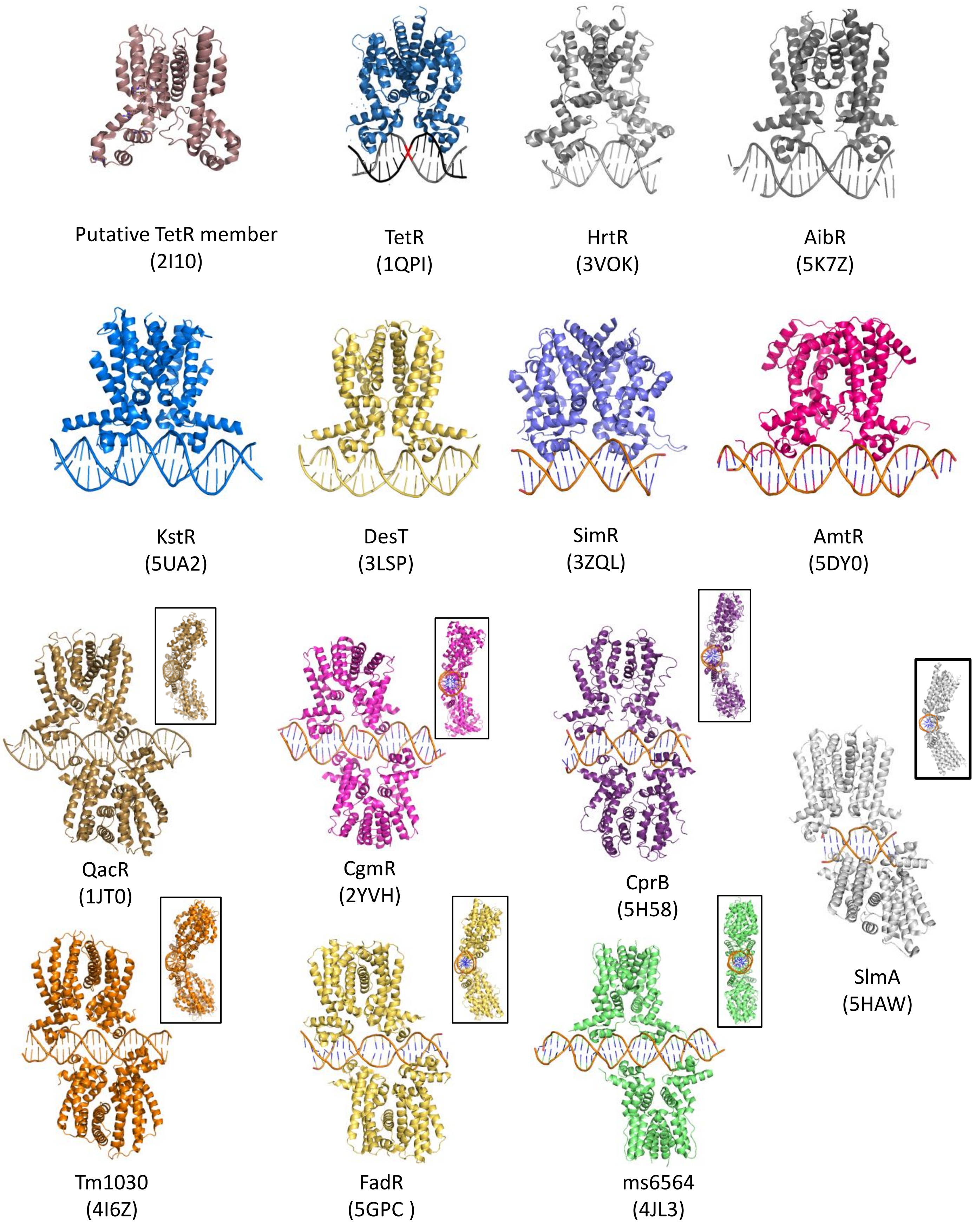


**Figure S3.** Structures of molecular-replacement search models. MR searches were performed with 14 TFR:DNA complexes and the apo-TFR structure with the highest sequence identity to CifR (PDB ID: 2I10). None of the MR searches yielded successful initial phase estimates for the $CifR_{TR}$:$dsDNA_{26L}$ complex dataset. Insets accompanying some of the models represent a side view (rotated by 90˚).

### Figure S4

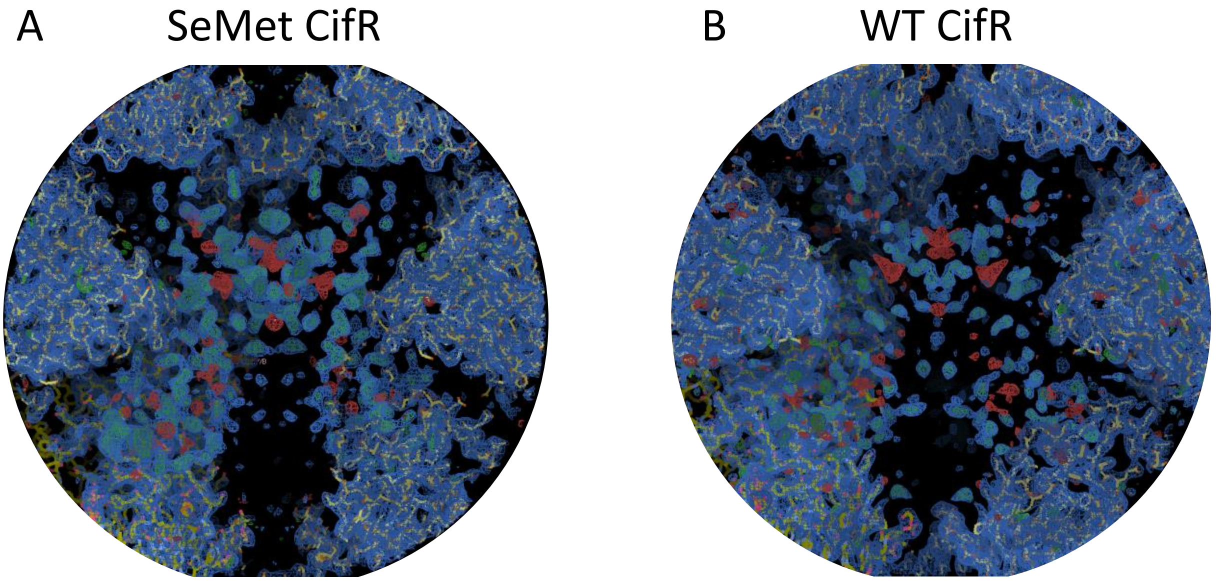


**Figure S4.** Electron density in the bulk solvent channel for apo-$CifR_{TR}$ in the SeMet dataset **(A)** and in the native dataset **(B)**. **(A, B)** Both 2m$F_O$-D$F_C$ and m$F_O$-D$F_C$ difference electron density maps are shown, contoured at 1.0σ (blue mesh) and ±3.0σ (positive, green mesh; negative, red mesh), respectively, together with the molecular structure corresponding to the $CifR_{TR}$:$dsDNA_{26L}$ complex (stick figures colored by atom type: C = yellow, N = blue, O = red, S = green).

### Figure S5

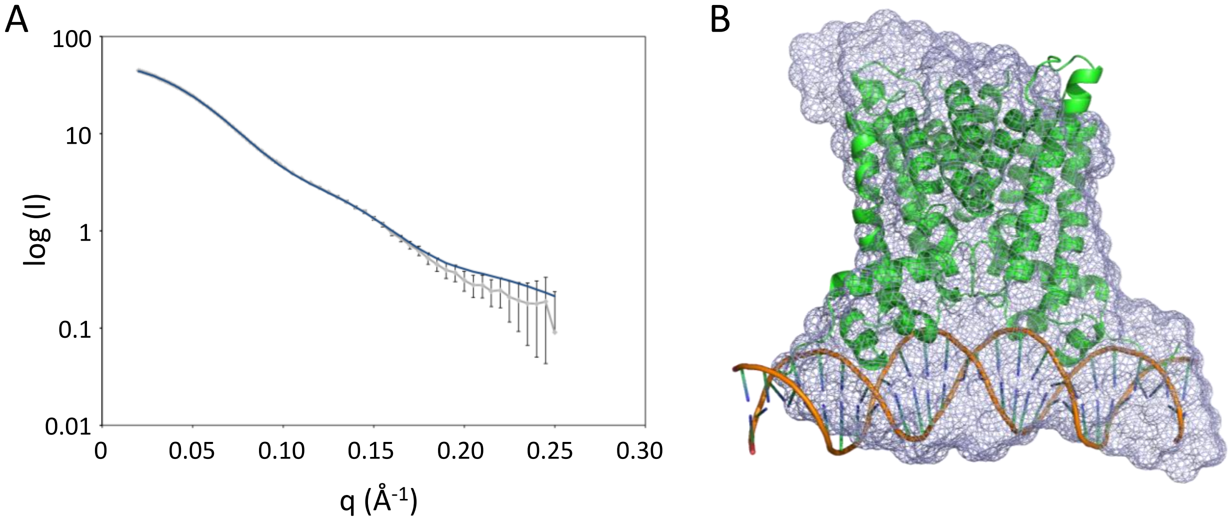


**Figure S5.** SAXS data for the CifR$_{TR}$:dsDNA$_{26L}$ complex are in good agreement with the crystal structure of the CifR$_{TR}$:dsDNA$_{26L}$ complex. **(A)** Theoretical SAXS data were calculated from the CifR$_{TR}$:dsDNA$_{26L}$ complex crystal structure (blue line) and fit to the experimental SAXS data. **(B)** A molecular envelope was calculated from the experimental SAXS data (blue mesh) and was used to dock the crystal structure of the CifR$_{TR}$:dsDNA$_{26L}$ complex (cartoon representation).

### Figure S6

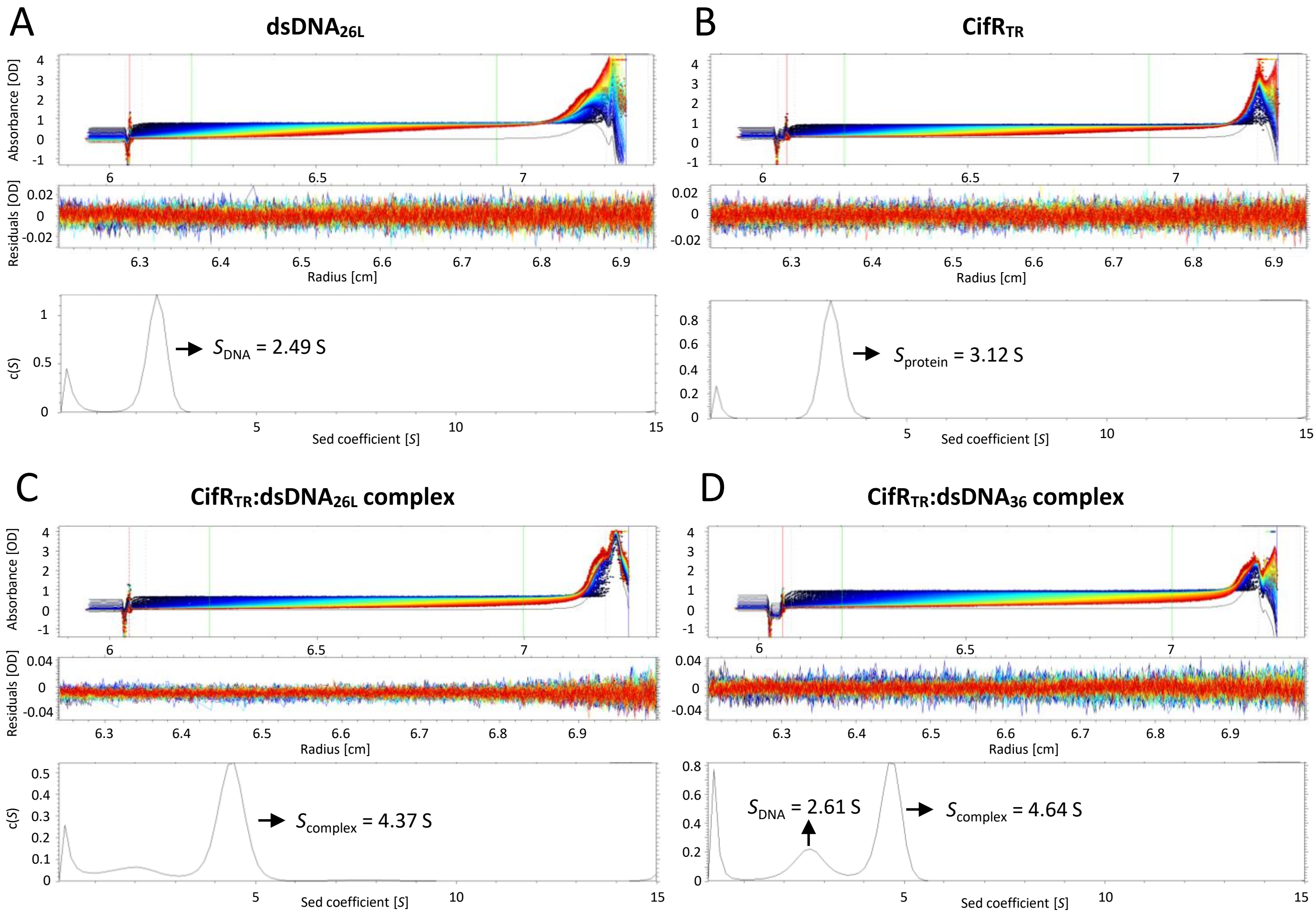


**Figure S6.** Velocity sedimentation analysis for $dsDNA_{26L}$ **(A)**, $CifR_{TR}$ **(B)**, $CifR_{TR}$:$dsDNA_{26L}$ **(C)**, and $CifR_{TR}$:$dsDNA_{36L}$ **(D)**. For each panel, the plot of the raw absorbance scans fitted with a nonlinear regression model (top), the plot of the fit residuals (middle), and the sedimentation coefficient distribution (bottom) are shown. In the plot of the raw absorbance scans, the red, blue, and green vertical lines indicate sample meniscus position, cell bottom position, and the fitting boundaries, respectively. (S = Svedberg unit, OD = optical density, c(*S*) = sedimentation coefficient concentration distribution.)

### Figure S7

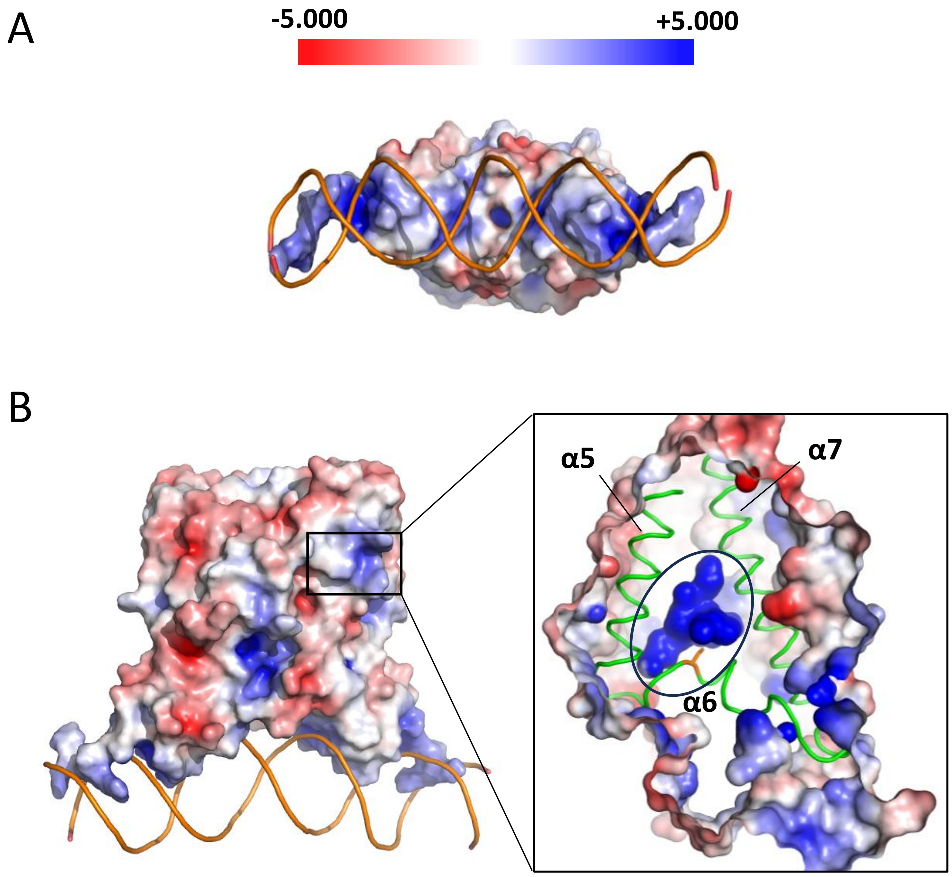


**Figure S7.** Surface electrostatic potential prediction for the $CifR_{TR}$ protein. The $CifR_{TR}$ model was extracted from the DNA-bound $CifR_{TR}$ complex structure and used for an APBS calculation. Red color indicates negative potential, while blue color indicates positive potential. **(A)** The bottom view of the DNA-binding surface of $CifR_{TR}$ shows an overall positive potential (blue; scale at top), complementing the net negative potential on the DNA backbone, with a concentration of positive potential towards the periphery of the binding site. **(B)** The front view of $CifR_{TR}$ shows a heterogeneous electrostatic profile. *Inset*: The potential surface of a cavity within the protein is shown on the right, highlighting a putative ligand-binding cavity (blue surface; marked by a black oval) with positive potential that is not visible from the outside and that is surrounded by helices α5-α7 (green ribbons). Cys107 is shown in orange (stick model).

### Figure S8

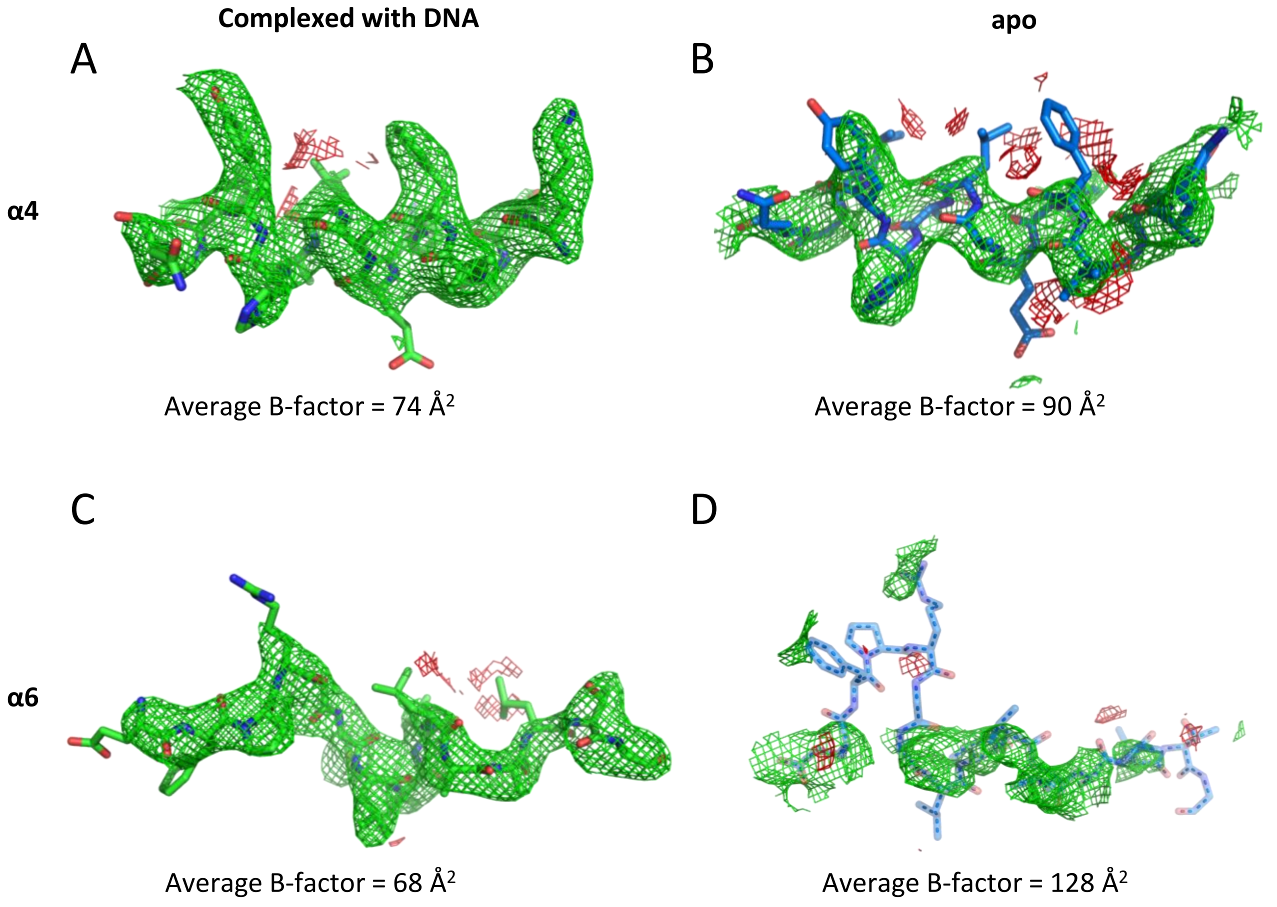


**Figure S8.** Polder OMIT maps of $\mathrm{CifR_{TR}}$:$\mathrm{dsDNA_{26L}}$ α4 **(A)**, apo-$\mathrm{CifR_{TR}}$ α4 **(B)**, $\mathrm{CifR_{TR}}$:$\mathrm{dsDNA_{26L}}$ α6 **(C)** and apo-$\mathrm{CifR_{TR}}$ α6 **(D)** helices. The $mF_O$-$DF_C$ electron density map and the model of the omitted residues 54-68 of chain A and chain M are shown for **(A)** and **(B)**, respectively. Omitted residues 102-116 of chain A and chain M are shown for **(C)** and **(D)**, respectively. The positive (green mesh) and negative (red mesh) electron density map peaks are contoured to ±3.0σ for **(A)** and **(C)** but were contoured to ±1.8σ in **(B)** and **(D)** to account for the 60% occupancy of chain M within the lattice. The model of the respective residues is shown in stick representation and colored by atom type. C= green, O=red, N=blue, S=yellow for **(A)** and **(C)**. C= light blue, O=red, N=blue, S=yellow for **(B)** and **(D)**.

### Figure S9

A

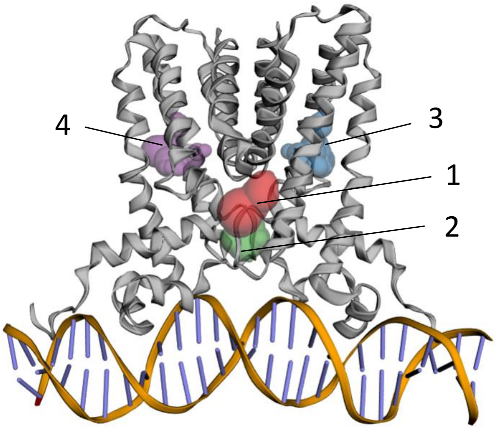


| PocID | Area (Å²) | Volume (Å³) |
|---|---|---|
| 1 | 103 | 76 |
| 2 | 134 | 67 |
| 3 | 184 | 53 |
| 4 | 160 | 52 |

B

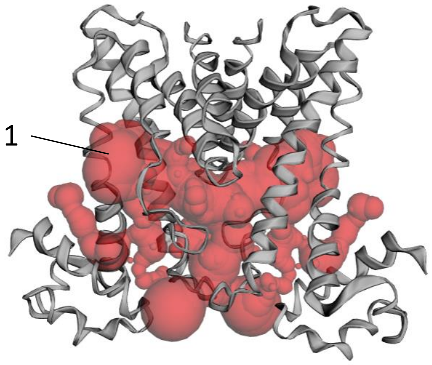


| PocID | Area (Å²) | Volume (Å³) |
|---|---|---|
| 1 | 2614 | 2133 |

**Figure S9.** Cavity distributions for CifR$_{TR}$. **(A)** Potential ligand-binding pockets (3 and 4) and entry and exit channels (1 and 2) were identified using CastP for CifR$_{TR}$:dsDNA$_{26L}$. **(B)** A potential ligand-binding pocket simulated using CastP for apo-CifR$_{TR}$. The area and volume for each pocket are shown in the table below each model.

### Figure S10

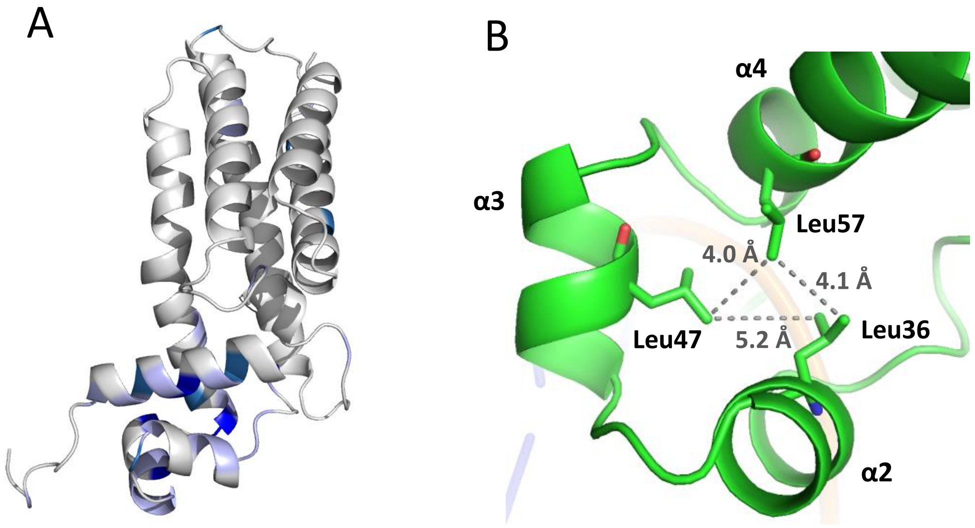


**Figure S10.** CifR$_{TR}$ sequence conservation and LBD:DBD packing interactions. **(A)** Mapping of the conserved residues in Figure S1 onto the CifR$_{TR}$-dsDNA$_{26L}$ complex structure (cartoon). Only one CifR monomer is shown for clarity. Residue coloring reflects sequence identity among the 938 TFRs analyzed: dark blue, >80%; sky blue, 60-80%; light grey blue, 40-60%; white, 0-40% identity. **(B)** Hydrophobic interactions (grey dashed lines) involving Leu36, Leu47 and Leu57 are shown.

### Figure S11

| TFRs | PDB id | #dimers/ operator | Operator sequence |
|---|---|---|---|
| QacR | 1JT0 | 2 | CTTATAGACCGATC \| GATCGGTCTATAAG<br>CTTATAGACCGA \| TCGATCGGTCTATAAG<br>CTTATAGACCGATCGA \| TCGGTCTATAAG |
| * CgmR | 2YVH | 2 | TAACTGTACCGACC \| GGTCGGTACAGTTA<br>TAACTGTACCGA \| CCGGTCGGTACAGTTA<br>TAACTGTACCGACCGG \| TCGGTACAGTTA |
| CprB | 5H58 | 2 | AGGCAGGCGGCACG \| GTCTGTTGAGTTC<br>AGGCAGGCGGCA \| CGGTCTGTTGAGTTC<br>AGGCAGGCGGCACGGT \| CTGTTGAGTTC |
| ms6564 | 4JL3 | 2 | TCATAAACGAGACGGTAC \| GTCTCGTCTTGTG<br>TCATAAACGAGACGG \| T \| ACGTCTCGTCTTGTG<br>TCATAAACGAGACGGTACGT \| C \| TCGTCTTGTG |
| Rv0078 | 6C31 | 2 | GTTACCGGCAG \| T \| CTGCTTGTAAA<br>GTTACCGGC \| A \| GTCTGCTTGTAAA<br>GTTACCGGCAGTC \| T \| GCTTGTAAA |
| *TM1030 | 4I6Z | 2 | GACTGACTGACA \| TGTCAGTCAGTC<br>GACTGACTGA \| CATGTCAGTCAGTC<br>GACTGACTGACATG \| TCAGTCAGTC |
| FadR | 5GPC | 2 | GATGAATGAAT \| ACTCATTCAT<br>GATGAATGA \| ATACTCATTCAT<br>GATGAATGAATAC \| TCATTCAT |
| SlmA | 5HAW | 2 | GTGAGT \| ACTCAC<br>GTGA \| GTACTCAC<br>GTGAGTAC \| TCAC |
| DarR | 8SVA | 2 | TAGATACTCC \| GGAGTATCTA<br>TAGATAC \| TCCGGAGTATCTA<br>TTGCTAC \| TCCGGAGTATCTA |

**Figure S11.** Operator DNA sequences of nine TFRs with two dimers binding to one operator DNA. Three rows of the same operator DNA are shown for each TFR. The *top row* shows the overall symmetry of entire operator. The *middle row* shows the symmetry pattern for the left embedded repeat. Except for DarR, the *bottom row* shows the symmetry pattern for the right embedded repeat. For DarR, the overall symmetry of entire operator (*top row*) aligns with one DarR dimer. The *middle row* shows the center for the other DarR dimer, which lacks an embedded repeat. The *bottom row* (surrounded by the dashed rectangle) shows the optimized operator sequence of the sequence engineered (substitutions in blue) to include an embedded repeat (59); no structure is yet available for this operator sequence. The bases with potential to form inverted repeats are underlined and in bold. The centers for two embedded repeats are highlighted in grey. Asterisks mark complexes for which only half of the operator DNA was used in the crystallization.

### Figure S12

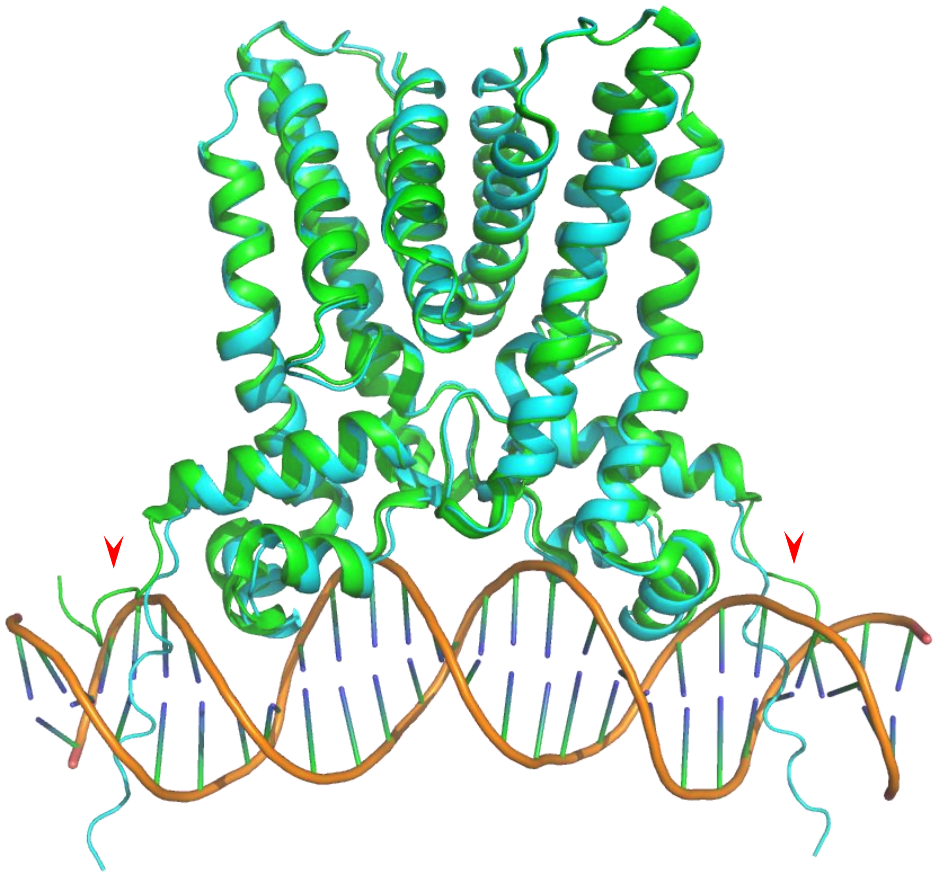

**Figure S12.** Least-squares superposition reveals excellent agreement (RMSD 0.6 Å) between the experimental $CifR_{TR}$:$dsDNA_{26L}$ dimer (green ribbon and orange helices; PDB 6NSN) and a computational model of the DNA-free conformation of CifR (cyan ribbon) generated by AlphaFold (62). The most significant divergence occurs at the N-terminal DNA-binding sequences found at the far left and far right of the figure (red arrows).

Figure S13

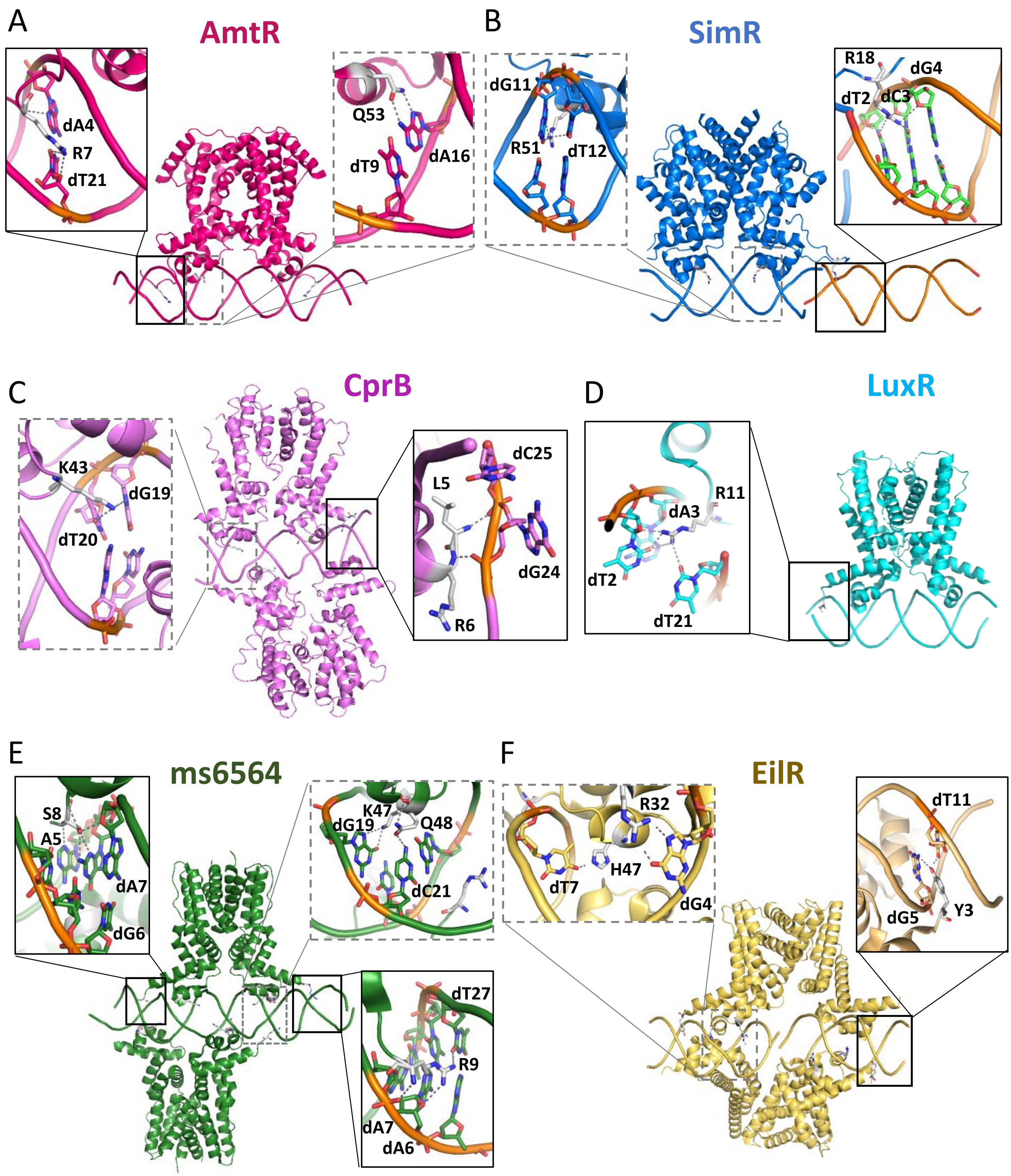


**Figure S13.** N-terminal interactions with the DNA minor groove. TFR:DNA complex structures are shown for AmtR [**A**; PDB entry 5DY0], SimR [**B**; 3ZQL], CprB [**C**; 5H58], LuxR [**D**; 7AMT], ms6564 [**E**; 4JL3], and EilR [**F**; 5VL9]. Main-chain structures (ribbons), base-interacting side chains (sticks), DNA backbones (traces), and interacting DNA bases (sticks) are shown. Protein-DNA interactions are magnified in the insets, with interacting side chains and DNA bases shown as stick figures, and the backbone of interacting bases highlighted in orange. Solid black and dashed grey boxes show major and minor groove interactions, respectively. Hydrogen bonds are shown as grey dashes.

### Table S1: *In vivo* primers and *in vitro* oligos used in this work.

| ***In vivo* primers** | **Sequence** |
|---|---|
| C99T_F | GCGAGACCGTCGCGCGCTTCACTTCCGATGAGTTCCCGCGCGG |
| C99T_R | CCGCGCGGGAACTCATCGGAAGTGAAGCGCGCGACGGTCTCGC |
| C107T_F | CCGATGAGTTCCCGCGCGGTACCCTGGTGGTGCTCGCGGCGCT |
| C107T_R | AGCGCCGCGAGCACCACCAGGGTACCGCGCGGGAACTCATCGG |
| C181R_F | AGGCCAAGGACCGGGTGCCTCGCGAGCGCCTGCTGGCGGTGGT |
| C181R_R | ACCACCGCCAGCAGGCGCTCGCGAGGCACCCGGTCCTTGGCCT |
| C107S_F | GATGAGTTCCCGCGCGGTAGCCTGGTGGTGCTCGCGGCG |
| C107S_R | CGCCGCGAGCACCACCAGGCTACCGCGCGGGAACTCATC |
| R6A_F | GCCCCATGGCAACGCGAGGCGCGCCACGGGCATTCGACAGGGA |
| R6A_R | TCCCTGTCGAATGCCCGTGGCGCGCCTCGCGTTGCCATGGGGC |
| CifR_delN10+3C_NdeI | GTCATATGCTCGAGGTGCTCTTCCAGGGCCCCGACAGGGACACCGCC |
| CifR_Cter_BamHI | CCGGATCCTCAGGGC |

| ***In vitro* oligos** | **Sequence** |
|---|---|
| *cifR-prox* $dsDNA_{36}$-up | CCTCCATTATTTGTATCGATCACTATAAATTTACTT |
| *cifR-prox* $dsDNA_{36}$-bottom | AAGTAAATTTATAGTGATCGATACAAATAATGGAGG |
| *cifR-prox* $dsDNA_{32+2nt}$-up | CCTCCATTATTTGTATCGATCACTATAAATTTAC |
| *cifR-prox* $dsDNA_{32+2nt}$-bottom | GGGTAAATTTATAGTGATCGATACAAATAATGGA |
| *cifR-prox* $dsDNA_{27}$-up | ATTATTTGTATCGATCACTATAAATTT |
| *cifR-prox* $dsDNA_{27}$-bottom | AAATTTATAGTGATCGATACAAATAAT |
| *cifR-prox* $dsDNA_{26L}$-up | TTATTTGTATCGATCACTATAAATTT |
| *cifR-prox* $dsDNA_{26L}$-bottom | AAATTTATAGTGATCGATACAAATAA |
| *cifR-prox* $dsDNA_{26R}$-up | ATTATTTGTATCGATCACTATAAATT |
| *cifR-prox* $dsDNA_{26R}$-bottom | AATTTATAGTGATCGATACAAATAAT |
| *cifR-prox* $dsDNA_{25}$-up | TTATTTGTATCGATCACTATAAATT |
| *cifR-prox* $dsDNA_{25}$-bottom | AATTTATAGTGATCGATACAAATAA |
| *cifR-prox* $dsDNA_{23+2nt}$-up | TTTATTTGTATCGATCACTATAAAT |
| *cifR-prox* $dsDNA_{23+2nt}$-bottom | AAATTTATAGTGATCGATACAAATA |
| *cifR-prox* $dsDNA_{23}$-up | TATTTGTATCGATCACTATAAAT |
| *cifR-prox* $dsDNA_{23}$-bottom | ATTTATAGTGATCGATACAAATA |
| *cifR-prox* $dsDNA_{20}$-up | TTTGTATCGATCACTATAAA |
| *cifR-prox* $dsDNA_{20}$-bottom | TTTATAGTGATCGATACAAA |
| *cifR-prox* $dsDNA_{26L}^{T4G}$-up | TTAGTTGTATCGATCACTATAACTTT |
| *cifR-prox* $dsDNA_{26L}^{T4G}$-bottom | AAAGTTATAGTGATCGATACAACTAA |
| *cifR-prox* $dsDNA_{26L}^{4\text{-}mix}$-up | TTATTTGTATCGATTCTGATAAATTT |
| *cifR-prox* $dsDNA_{26L}^{4\text{-}mix}$-bottom | AAATTTATCAGAATCGATACAAATAA |
| *morB-prox* $dsDNA_{26L}$-up | ATATCTGTATCGGTCGCTAAATAATA |
| *morB-prox* $dsDNA_{26L}$-down | TATTATTTAGCGACCGATACAGATAT |

### Table S2: Crystallization conditions for initial screening hits and final conditions for data collection

| Complex | $CifR_{RT}$-$dsDNA_{36}$ | $CifR_{RT}$-$dsDNA_{27}$ | $CifR_{RT}$-$dsDNA_{26L}^{MSE}$ | $CifR_{RT}$-$dsDNA_{26L}$ |
|---|---|---|---|---|
| **Initial condition** | 16% [*w/v*] PEG3350<br>0.011 M nickel chloride<br>0.015 M magnesium chloride | 20% [*w/v*] glycerol<br>14.4% [*w/v*] PEG8000<br>0.16 M calcium acetate<br>0.08 M sodium cacodylate, pH6.5 | 0.2 M magnesium chloride<br>10% [w/v] PEG1000<br>10% [w/v] PEG8000<br>0.1 M Tris, pH 7.5 | 15% [*w/v*] PEG4000<br>0.2 M magnesium chloride<br>0.1 M Tris, pH 8.5 |
| **Final condition** | N/A | N/A | 0.2 M magnesium chloride<br>10% [w/v] PEG1000<br>10% [w/v] PEG8000<br>0.1 M Tris, pH 7.5 | 16% [w/v] PEG4000<br>0.2 M magnesium chloride<br>0.1 M Tris, pH 8.3 |
